# Dysregulation of astrocyte-secreted pleiotrophin contributes to neuronal structural and functional deficits in Down Syndrome

**DOI:** 10.1101/2023.09.26.559633

**Authors:** Ashley N. Brandebura, Quinn N. Asbell, Mariel Kristine B. Micael, Nicola J. Allen

**Affiliations:** The Salk Institute for Biological Studies, Molecular Neurobiology Laboratory, 10010 North Torrey Pines Road, La Jolla, CA 92037, USA; University of California San Diego, Department of Psychology, Muir Ln, La Jolla, CA 92093, USA

**Keywords:** Astrocyte, dendritic spines, synapse, pleiotrophin, dendrite, neuron, Down Syndrome, neurodevelopmental disorder, trisomy 21, visual cortex

## Abstract

Neuronal dendrite patterning and synapse formation are tightly regulated during development to promote proper connectivity. Astrocyte-secreted proteins act as guidance and pro-synaptogenic factors during development, but little is known about how astrocytes may contribute to neurodevelopmental disorders. Here we identify down-regulation of the astrocyte-secreted molecule pleiotrophin as a major contributor to neuronal morphological alterations in the Ts65Dn mouse model of Down Syndrome. We find overlapping deficits in neuronal dendrites, spines and intracortical synapses in Ts65Dn mutant and pleiotrophin knockout mice. By targeting pleiotrophin overexpression to astrocytes in adult Ts65Dn mutant mice *in vivo*, we show that pleiotrophin can rescue dendrite morphology and spine density and increase excitatory synapse number. We further demonstrate functional improvements in behavior. Our findings identify pleiotrophin as a molecule that can be used in Down Syndrome to promote proper circuit connectivity, importantly at later stages of development after typical periods of circuit refinement have completed.

## Introduction

Down Syndrome (DS) is the most common form of genetic intellectual disability, caused by trisomy of human chromosome 21 (HSA21) and affecting 1 in 700 live births^1^. Individuals with DS have neurodevelopmental changes that result in hyperactivity, as well as altered information processing and memory impediments^2^. Histological studies have identified changes to neuronal morphology during development in DS which may be linked to these functional impairments. For example, Layer 5 pyramidal neurons in the visual cortex have shorter and less complex dendrites than those from neurotypical individuals^3^. Additionally, decreased dendritic spine density is reported in pyramidal neurons of the motor cortex, visual cortex and hippocampus^4–6^.

These classical studies were performed decades ago, but the field is still in search of therapies that can improve alterations to neuronal morphology, synaptic circuitry and cognitive function. The development of complex neuronal morphology and formation of synapses is not only regulated in a cell-autonomous manner. Rather, astrocytes, an abundant type of glial cell in the brain, play essential roles in regulating neuronal synapse number and function during development^7^. One key mechanism by which astrocytes modulate synapses is through the secretion of proteins that guide multiple stages of synaptogenesis, including synapse formation and maturation^8–11^. With respect to DS, there is accumulating evidence for astrocyte involvement. Studies have demonstrated that wildtype (WT) neurons, cultured *in vitro* with DS astrocytes or with DS astrocyte conditioned media (ACM), acquired DS phenotypes such as reduced spine and synapse number and retention of immature filopodia spines^12,13^. One *in vitro* study showed that addition of Thrombospondin1 (Tsp1), a protein with known roles in synaptogenesis^14^, to the media in WT neuron/DS astrocyte co-cultures could partially recover the spine deficits induced by DS astrocytes^12^. However, the incomplete rescue with Tsp1 suggests there are additional astrocyte-secreted proteins contributing to these deficits. A second study showed that DS ACM inhibits the functional maturation of WT neuron voltage-gated sodium and potassium channels, which are necessary for action potential propagation, whereas adding WT ACM to DS neurons improved ion channel conductance in the trisomic neurons^15^. Together these studies demonstrate that neuronal phenotypes observed in DS are not solely intrinsic to the neuron and are heavily influenced by the astrocyte genotype, and in particular by the balance of astrocyte-secreted proteins.

To identify how astrocyte protein secretion is altered in DS, we performed unbiased mass spectrometry proteomics analysis of the ACM from cultured primary mouse cortical astrocytes isolated from the Ts65Dn mouse model of DS^16^. This study revealed that astrocyte protein secretion in DS is highly dysregulated, suggesting that altered secretion of multiple proteins from astrocytes may have detrimental effects on neuronal circuit formation in DS. However, the functional impact of these protein changes on neuronal dendrite morphology and synaptic circuitry in DS is not known. Additionally, prior studies were performed *in vitro*, so it is not known if modulation of astrocyte-secreted protein levels can restore aberrant neuronal circuitry *in vivo*. The goal of the present study is to identify a specific astrocyte-secreted protein that can be used to correct neuronal dendritic and synaptic deficits in neurodevelopmental disorders *in vivo*, focusing on DS. We identified the astrocyte-secreted heparin-binding growth factor pleiotrophin (Ptn) as a top candidate. Ptn is among the top ten down-regulated proteins secreted from Ts65Dn astrocytes^16^ and has a strong enrichment in cortical astrocytes compared to other cell types^17^. Prior *in vitro* work showed that Ptn is a neurite outgrowth inducing factor via interactions with heparan sulfate side chains and chondroitin sulfate proteoglycans in the extracellular matrix^18^, and that Ptn is a neurite outgrowth factor *in vivo* after spinal cord injury^19^. Ptn has also been demonstrated to have neuroprotective functions in the context of other acute injuries such as cocaine-induced cytotoxicity^20^, but its roles in the context of neurodevelopmental disorders such as Down Syndrome are not known. Furthermore, knockdown of Ptn in neural stem cells in the dentate gyrus results in decreased dendritic complexity and spine density of newly born hippocampal neurons, demonstrating that in the healthy brain Ptn can regulate neuronal development^21^.

Based on the ability of Ptn to promote neuronal development in the healthy brain, we hypothesized that downregulation of Ptn in astrocytes may be contributing to observed phenotypes of stunted dendrite growth and reduced dendritic spine density in DS. To address this, we explore the non-cell autonomous regulation of neuronal dendrite morphology and synaptic circuitry by Ptn in both neurotypical development and in DS. We use the visual cortex as a model system due to its well-characterized circuitry and previously described phenotypes of shorter and less complex dendrites in pyramidal neurons in DS^3,6^. We identify that genetic loss of Ptn results in neuronal dendrite morphological alterations and synaptic phenotypes that are highly overlapping with those observed in DS. We further demonstrate that viral-mediated targeting of Ptn overexpression specifically to astrocytes in adult Ts65Dn mice ameliorates many aspects of the aberrant neuronal morphology and connectivity *in vivo*, as well as improves cognitive function in two separate memory tasks. Importantly, these results are achieved with Ptn delivery at adult stages after aberrant circuitry has developed, indicating that Ptn can reverse impairments after developmental windows have closed. In future these findings may be translatable to other neurodevelopmental disorders with similar circuit aberrations.

## Results

### *Ptn* expression is developmentally regulated and is decreased in Ts65Dn astrocytes *in vivo*

In the visual cortex (VC) dendrite outgrowth is prominent from birth^22^, synapse formation is high from P7^23,24^, synaptic maturation begins ∼P14^25^ and circuit stabilization is achieved by ∼P120 (Figure 1A). In neurotypical development *Ptn* has widespread expression throughout the brain, and prominent expression in upper cortical layers (Figure S1A-A’). To ask how *Ptn* expression correlates with important periods of synaptic development, we utilized a previously published astrocyte RiboTag RNA-sequencing dataset of the VC^26^ to determine the temporal profile of *Ptn* expression, which shows peak levels of *Ptn* during periods of synaptogenesis and synaptic maturation (Figure 1B). We validated the expression profile of *Ptn* using single molecule fluorescent in situ hybridization (smFISH) in Aldh1L1-EGFP mice at P7, P14 and P30, where astrocytes are labeled with EGFP (Figure 1C-E). The upper cortical layers were chosen for quantification based on a previous report that *Ptn* is an upper layer astrocyte marker^26^. We found that *Ptn* has highest expression at P14 (Layer 1: 30.9 ± 3.8 a.u.; Layer 2/3: 40.2 ± 4.4 a.u.) and significantly decreases by P30 (Layer 1: 18.8 ± 2.1 a.u.; Layer 2/3: 21.1 ± 3.2 a.u.; Figure 1F-G).

**Figure 1.**
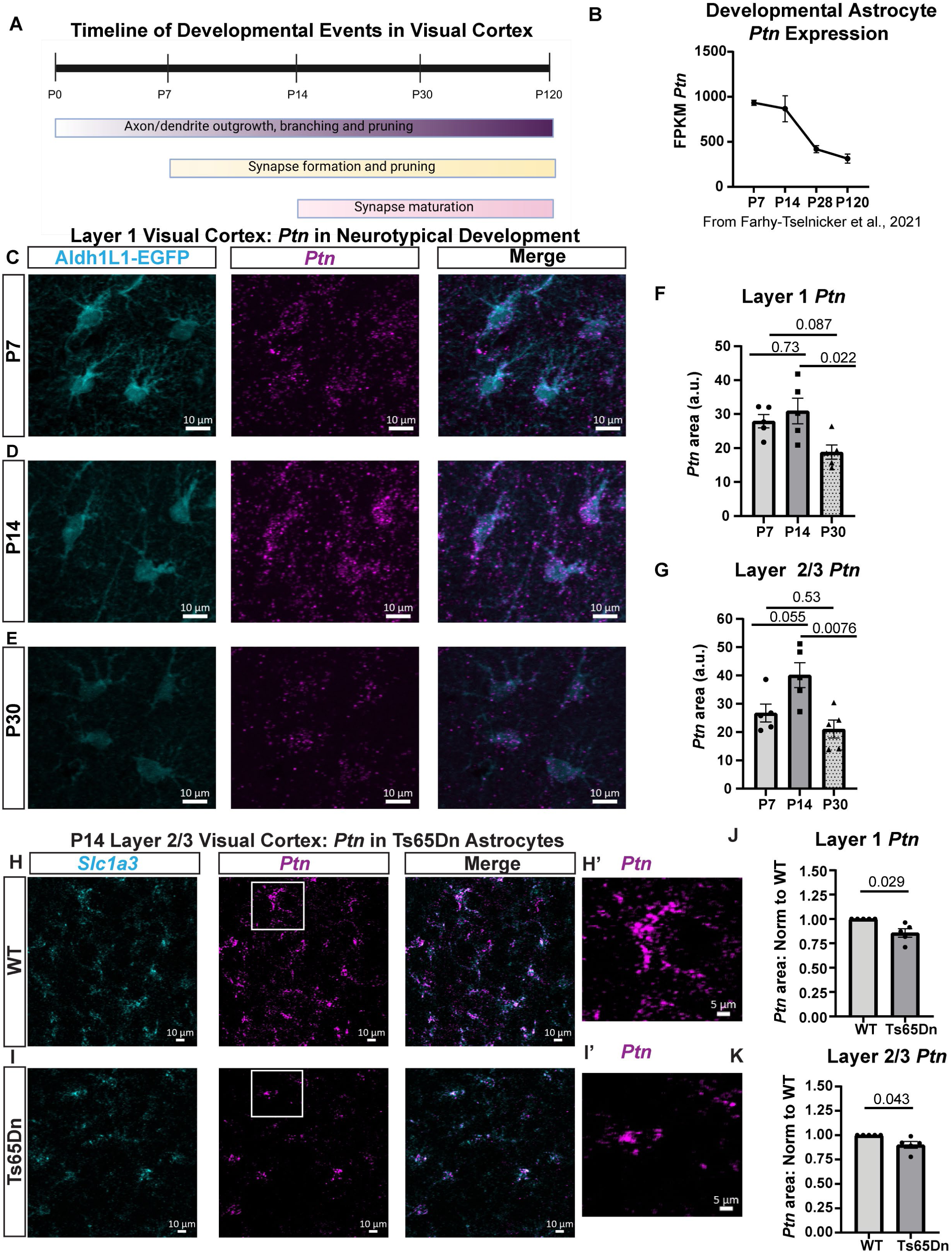
Developmental expression of astrocyte *Ptn* mirrors the timeframe for formation and maturation of cortical synapses and *Ptn* is reduced in Ts65Dn Mut astrocytes. (A) Timeline of developmental events in visual cortex (VC). (B) Astrocyte RiboTag RNA-Sequencing data from VC shows peak *Ptn* expression in astrocytes corresponding to the timeframe for synapse formation and maturation. Data from Farhy-Tselnicker et al., 2021. (C-E) Representative images of *Ptn* mRNA (magenta) in Aldh1L1-EGFP+ astrocytes (cyan) at P7, P14 and P30 in VC Layer 1. Scale bars = 10 um. (F-G) Quantification of *Ptn* area (a.u.) in astrocytes across development in Layer 1 and Layer 2/3. Statistics by one-way ANOVA with Tukey’s. (H-I) Example images of *Ptn* (magenta) in *Slc1a3*+ astrocytes (cyan) at P14 in VC Layer 2/3 of WT and Ts65Dn Mut mice. Scale bars = 10 um. (H’-I’) Zoomed images of *Ptn* within WT and Ts65Dn Mut astrocytes. Scale bars = 5 um. (J-K) Quantification of average *Ptn* area within Ts65Dn Mut astrocytes normalized to WT in Layer 1 and Layer 2/3. Statistics by one-sample t-test. N = 5 mice/timepoint in C-G and N = 5 mice/group in H-K.

We previously found that cultured Ts65Dn astrocytes have decreased *Ptn* mRNA and decreased secreted Ptn protein compared to wildtype astrocytes (Figure S1B-C; RNA-Seq: 173.8 ± 8.4 FPKM in WT, 114.9 ± 16.7 in Ts65Dn; Mass Spectrometry: 0.50 ± 0.13 NSAF in WT, 0.088 ± 0.034 NSAF in Ts65Dn ACM). Based on these findings, we investigated using smFISH whether *Ptn* expression is reduced in DS astrocytes *in vivo*. *Ptn* expression was quantified within *Slc1a3*+ astrocytes from Ts65Dn WT and Mutant (Mut) mice at P14 (Figure 1H-I; 1H’-I’). Ts65Dn Mut astrocytes have a significant reduction in *Ptn* in Layer 1 and Layer 2/3 astrocytes (Layer 1: 0.86 ± 0.043-fold, Layer 2/3: 0.90 ± 0.034-fold; Figure 1J-K). Altogether these data demonstrate that *Ptn* expression in astrocytes is developmentally regulated to coincide with peak synapse formation and maturation in the VC, and that *Ptn* is decreased in Ts65Dn Mut astrocytes *in vivo*, suggesting it may contribute to neuronal and synaptic phenotypes in DS.

### Astrocyte-secreted pleiotrophin promotes dendrite outgrowth and branching

Given that Layer 5 pyramidal neurons have dendritic outgrowth and branching impairments in humans with DS^3,6^ and that pyramidal neuron dendrites project to upper layers where *Ptn* is highly expressed^26^, we asked if Ptn can induce dendrite outgrowth in pyramidal neurons *in vitro.* This was approached by culturing cortical astrocytes from WT or a previously characterized Ptn knockout (KO) mouse model^27^, validated using smFISH that Ptn KO mice do not have detectable *Ptn* (Figure S2A). We utilized a cell culture model where WT cortical neurons are grown in minimal media (MM; without growth factors added), astrocyte conditioned media (ACM) from WT astrocytes (WT ACM), ACM from Ptn KO astrocytes (Ptn KO ACM), or Ptn KO ACM + recombinant Ptn (Rescue ACM; Figure 2A). We added recombinant Ptn at 0.025 ug/well after determining a physiological range of Ptn secretion from WT cultured astrocytes (Figure S2B). Dendrites were labeled with Map2, combined with nuclear Ctip2 (arrowheads in Figure 2B) to selectively enrich for Layer 5 pyramidal neurons^28^. Neurons cultured with WT ACM showed enhanced dendrite outgrowth compared to minimal media (average of 155.6 ± 9.1 um in MM, 225.7 ± 19.7 um in WT ACM). However, neurons cultured with Ptn KO ACM had significantly stunted dendrite outgrowth (115.8 ± 10.1 um). When neurons were exposed to Ptn KO ACM + recombinant Ptn protein, the total dendrite length was partially recovered (180.1 ± 13.2 um; Figure 2C; Figure S2C). We further used Sholl analysis to determine if there were significant differences in branching patterns as a measure of dendritic complexity. The number of Sholl intersections was graphed as a function of distance from the cell body (Figure 2D) and the area under the curves was quantified (Figure 2E). We observed that WT ACM induced more complex branching of dendrites (area of 121.7 ± 7.8 in MM, 181.3 ± 16.2 in WT ACM), whereas Ptn KO ACM impaired dendrite branching (90.4 ± 8.1), and Ptn KO ACM + recombinant Ptn protein was able to partially recover branching (144.1 ± 11.0). These experiments identify that astrocyte-secreted Ptn is necessary and sufficient to regulate neuronal dendrite outgrowth and branching *in vitro*.

**Figure 2.**
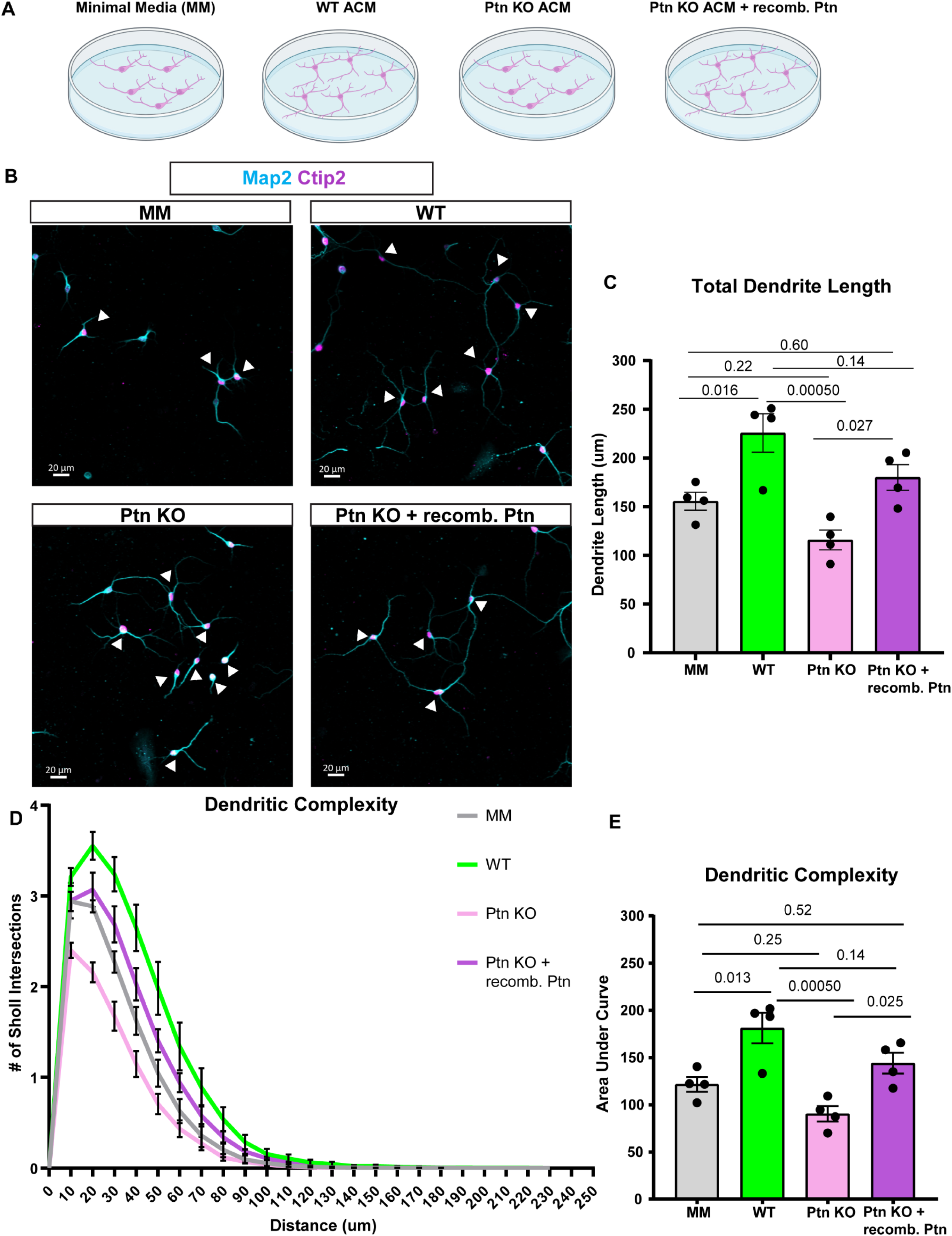
Genetic loss of Ptn from astrocytes impairs dendrite outgrowth and dendritic complexity *in vitro*. (A) Schematic demonstrating the *in vitro* experimental design to determine if astrocyte-derived Ptn modulates dendrite outgrowth and branching in WT cortical neurons. (B) Example images of each culture condition (MM: top left; WT ACM: top right; Ptn KO ACM: bottom left; Ptn KO ACM + recombinant Ptn: bottom right) showing WT pyramidal neurons labeled by Ctip2 (magenta; arrowheads) and their dendrites labeled by Map2 (cyan). Scale bars = 20 um. (C) Quantification of average total dendrite length for each condition shows that WT ACM induces dendrite outgrowth, whereas Ptn KO ACM fails to induce outgrowth above minimal media. Addition of recombinant Ptn (0.025 ug) to Ptn KO ACM partially restores dendrite outgrowth. Statistics by one-way ANOVA with Tukey’s. (D) Sholl curves of each culture condition representing the number of dendrite branches at 10 um increments from the cell body. (E) Quantification of area under the curve for Sholl graphs. WT ACM induces dendrite branching whereas Ptn KO ACM impairs branching and addition of Ptn protein to KO ACM partially restores branching. Statistics by one-way ANOVA with Tukey’s. N = 4 independent experiments.

### Pleiotrophin KO mice phenocopy DS Ts65Dn Mut mice in dendrite outgrowth, dendrite branching and spine density alterations *in vivo*

Knowing that Ptn can induce neuronal dendrite outgrowth and branching *in vitro*, we next assessed whether Ptn KO mice have *in vivo* phenotypes similar to those reported in humans with DS, and whether these phenotypes are also present in Ts65Dn mice. To do this we crossed the Ptn KO and Ts65Dn mice to a Thy1-YFP reporter line that sparsely labels Layer 5 pyramidal neurons^29^. Total dendrite length, dendritic complexity and spine density of neurons in the VC were analyzed at P30 and P120 to represent developmental and mature timepoints, respectively. We found that Layer 5 pyramidal neurons of Ts65Dn Mut mice have significantly shorter total dendrite length at P30 (average of 718.9 ± 108.4 um in WT, 432.5 ± 53.6 um in Mut, Figure A-C) and at P120 (644.7 ± 35.3 um in WT, 519.5 ± 42.4 um in Mut; Figure 3C). Sholl analysis showed decreased dendritic complexity at both ages when area under the curve was quantified (P30: area of 537.4 ± 75.7 in WT, 305.1 ± 46.9 in Mut; P120: 454.2 ± 31.2 in WT, 333.7 ± 34.2 in Mut; Figure 3D). This validates that Ts65Dn Mut mice accurately reflect the human phenotypes observed in DS of decreased dendritic arbor length and complexity. We next performed the same quantifications in the Ptn x Thy1-YFP line. We observed that Ptn KO mice have shorter total dendrite length at P30 (average of 635.7 ± 81.2 um in WT, 389.0 ± 57.8 µm in KO; Figure 3E-G), but interestingly the dendrite length recovers by P120 (506.5 ± 74.5 um in WT, 535.1 ± 75.5 um in KO; Figure 3G). Similarly, Sholl analysis revealed decreased dendritic complexity in KO mice at P30 (area of 506.7 ± 61.4 um in WT, 310.8 ± 48.2 µm in KO) that recovered by P120 (367.0 ± 46.5 in WT, 391.7 ± 47.7 in KO; Figure 3H). Distributions for individual dendrite lengths and Sholl graphs are shown in the associated supplemental figure (Figure S3A-D).

**Figure 3.**
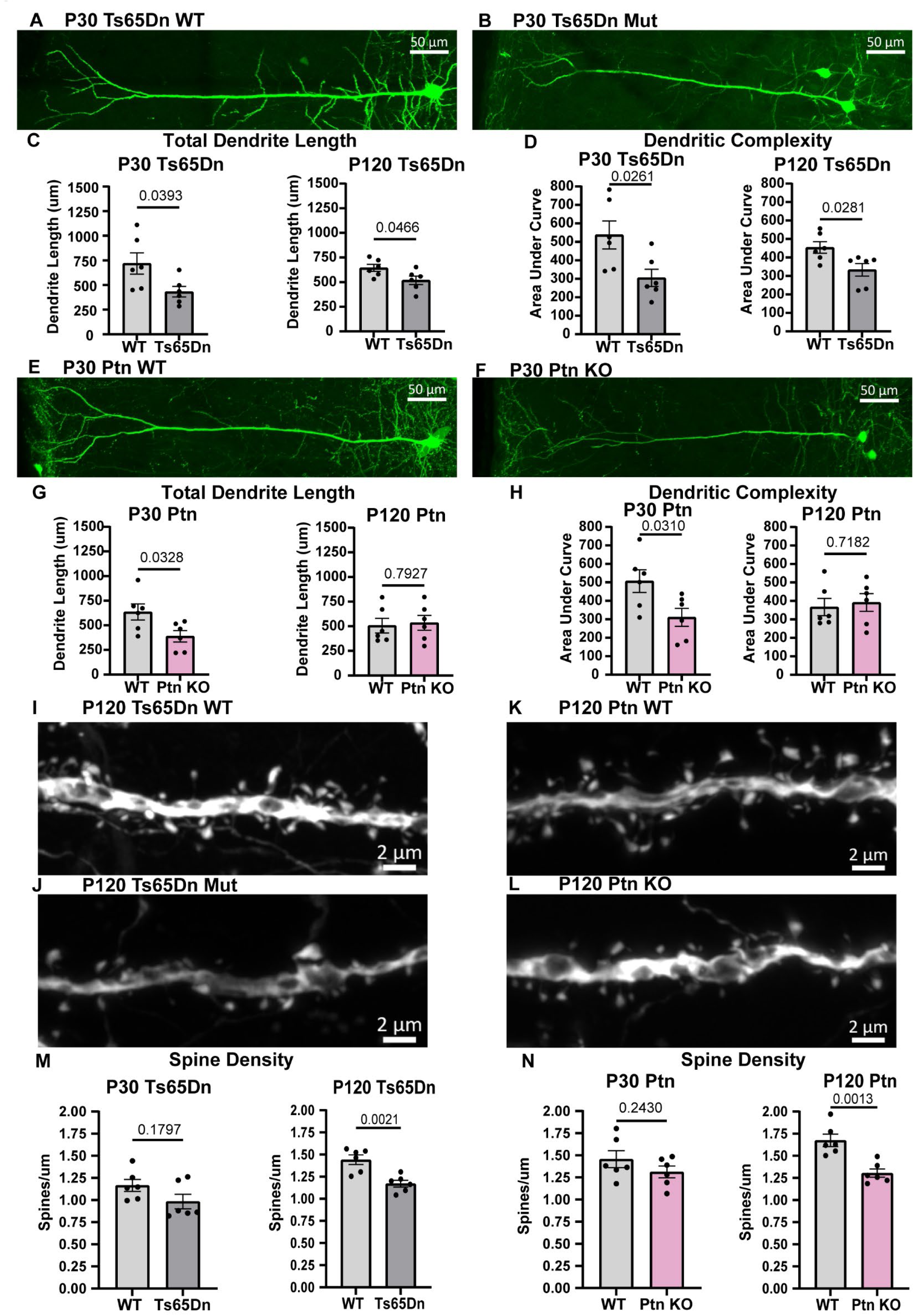
Ptn KO and Ts65Dn Mut mice display parallel dendrite and spine phenotypes. (A-B) Example images of Ts65Dn WT and Mut Layer 5 pyramidal neurons at P30. Scale bars = 50 um. (C) Data showing total dendrite length is reduced in Ts65Dn Mut mice at P30 and P120. Quantification of area under the curve from Sholl analysis shows that Ts65Dn Mut mice have less complex branching patterns than WT mice at both ages. (E-F) Example images of dendrites in Ptn WT and KO mice at P30. Scale bars = 50 um. (G) Total dendrite length is reduced in Ptn KO mice at P30 but recovers at P120. (H) Quantification of area under the curve for Sholl analysis shows that Ptn KO mice have reduced dendritic complexity at P30 that recovers by P120. (I-L) Example images of dendritic spines in Ts65Dn WT and Mut mice (I-J) and Ptn WT and KO mice (K-L) at P120, imaged in Layers 1-3. Scale bars = 2 um. (M-N) Quantification of average spine density shows Ts65Dn Mut mice have reduced spine density at P120 (M) and Ptn KO mice have reduced spine density at P120 (N). N = 6 mice/group. All statistics by unpaired t-test except P120 Ts65Dn AUC (D) and P30 Ts65Dn spine density (M) with Mann-Whitney.

As spine number and morphology are altered in DS, we quantified spine density in Ptn KO and Ts65Dn mice, focusing on dendrites from Layer 5 neurons that had their primary branching point in upper layers. Ts65Dn Mut mice had a significantly decreased spine density at P120 (average of 1.44 ± 0.054 spines/um in WT, 1.17 ± 0.038 spines/um in Mut; Figure 3I-J and M), but no statistically significant difference at P30 (average of 1.17 ± 0.068 spines/um in WT, 0.98 ± 0.083 spines/um in Mut; Figure 3M). Ptn KO mice also had reduced spine density at P120 (average of 1.67 ± 0.070 spines/um in WT, 1.30 ± 0.047 spines/um in KO; Figure 3K-L and N) but not at P30 (average of 1.46 ± 0.096 spines/um in WT, 1.31 ± 0.066 spines/um in KO; Figure 3N). Data from individual dendrites are shown in the associated supplemental figure (Figure S3E and G). Spine morphology was also assessed using previously published criteria for immature filopodia, thin, and long thin spines as well as more mature stubby and mushroom morphologies^30,31^. We observed an increased percentage of filopodia spines at P120 in Ts65Dn Mut mice (average of 11.5 ± 1.0% of total spines in WT, 20.2 ± 2.1% of total spines in Mut; Figure S3F), with Ptn KO mice showing the same trend (Figure S3H). The parallel phenotypes observed in Ptn KO and Ts65Dn Mut mice demonstrate that Ptn contributes to dendrite growth as well as spine formation and/or maturation during neurotypical development, and suggest that decreased Ptn in Ts65Dn Mut astrocytes may contribute to neuronal phenotypes.

### Overlapping synaptic alterations are observed in Pleiotrophin KO and DS Ts65Dn Mut mice

*Ptn* expression peaks during a robust period of synapse formation and maturation (Figure 1), so we hypothesized it may be involved in regulating excitatory synapse number. We assessed whether Ts65Dn Mut and Ptn KO mice have altered numbers of excitatory synapses compared to their WT counterparts. We used antibodies against the vesicular glutamate transporter 1 (Vglut1) and the GluA2 AMPA receptor subunit to label pre- and post-synaptic puncta of intracortical synapses, imaged in Layer 2/3. Ts65Dn Mut mice have a significant decrease in colocalized puncta (Figure 4A) at P30 (0.89 ± 0.028-fold compared to WT; Figure 4B) and this phenotype remains at P120 (0.82 ± 0.062-fold compared to WT; Figure 4C-D). Similarly, we found significantly decreased colocalized puncta (Figure 4E) at P30 in Ptn KO mice (0.83 ± 0.049-fold compared to WT), driven by a decrease in GluA2 puncta (0.92 ± 0.026-fold compared to WT) with no effect on Vglut1 puncta (0.97 ± 0.022-fold compared to WT; Figure 4F). These impairments remained at the P120 timepoint (Colocalized: 0.83 ± 0.043-fold, GluA2: 0.94 ± 0.019-fold, Vglut1: 0.96 ± 0.026-fold; Figure 4G-H).

**Figure 4.**
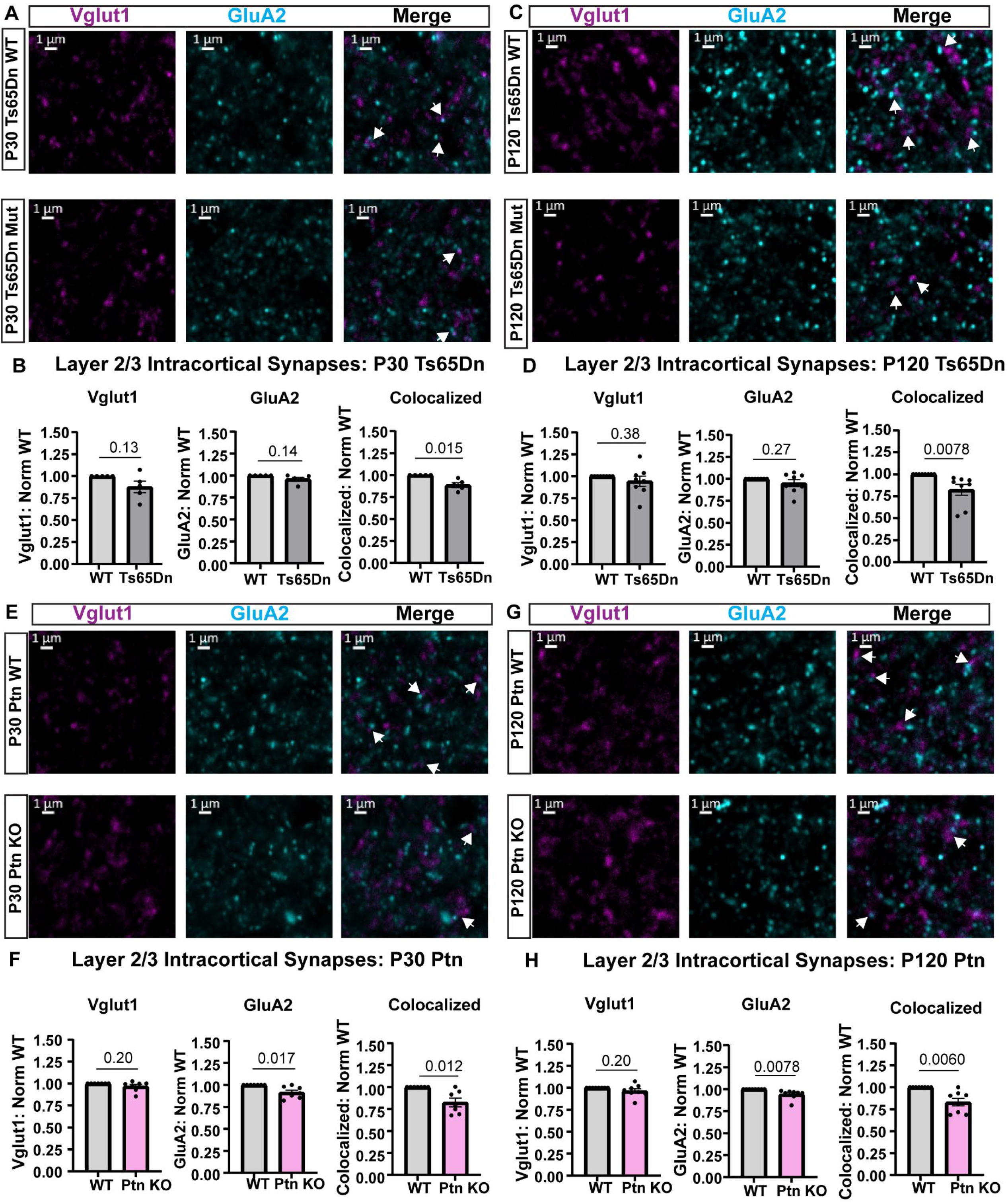
Ts65Dn Mut and Ptn KO mice have similar phenotypes in intracortical excitatory synapses. (A) Example images of Vglut1 (magenta) and GluA2 (cyan) labeling pre- and postsynaptic puncta, respectively, and their colocalization (white arrows) in the merged images at P30 in Ts65Dn WT and Mut mice, in Layer 2/3 of VC. (B) Quantification at P30 shows a significant decrease in colocalized puncta. (C) Example images as in (A) at P120 in Ts65Dn WT and Mut mice. (D) Quantification shows Ts65Dn Mut mice have decreased colocalized puncta at P120. Example images of Vglut1 (magenta) and GluA2 (cyan) and their colocalization (white arrows) at P30 in Ptn WT and KO mice. (F) Ptn KO mice have a significant decrease in post-synaptic GluA2 puncta and colocalized puncta at P30. (G) Representative images as in (E) at P120 in Ptn WT and KO mice. (H) Quantification at P120 shows the decreased GluA2 and colocalized puncta phenotypes remain at this age. Scale bars for all images = 1 um. N = 6-8 mice/group. Statistics by one-sample t-test for all except P120 Ts65Dn Colocalized Puncta and P120 Ptn KO GluA2 puncta assessed by one-sample Wilcoxon Signed-Rank test.

We also examined thalamocortical synapses using Vglut2 with GluA2 colocalization in Layer 1. Interestingly, there were clear distinctions between the Ts65Dn Mut and Ptn KO mice in these analyses. Ts65Dn Mut mice did not show any deficits in GluA2, Vglut2, or colocalized puncta (GluA2: 1.022 ± 0.014-fold, Vglut2: 1.015 ± 0.035-fold, Colocalized: 1.024 ± 0.029-fold; Figure S4A-B) at P30 or at P120 (GluA2: 1.068 ± 0.050-fold, VGlut2: 0.96 ± 0.055-fold, Colocalized: 0.99 ± 0.074-fold; Figure S4C-D). On the other hand, at P30 Ptn KO mice had decreased GluA2 (0.91 ± 0.025-fold), decreased Vglut2 (0.92 ± 0.021-fold) and decreased colocalized puncta (0.87 ± 0.046-fold; Figure 4SE-F). These differences remained at P120 (GluA2: 0.91 ± 0.028-fold, Vglut2: 0.90 ± 0.041-fold, Colocalized: 0.82 ± 0.044-fold; Figure S4G-H). The collective evidence demonstrates that both Ts65Dn Mut and Ptn KO mice have impaired intracortical synapse number in Layer 2/3 that persists to adulthood, while only Ptn KO mice have impaired thalamocortical synapses. This suggests roles of Ptn in thalamocortical synapse development that are compensated for in Ts65Dn Mut mice.

### Astrocyte-targeted pleiotrophin overexpression rescues dendrite and spine phenotypes in DS Ts65Dn Mut mice

The overlapping neuronal alterations observed in Ts65Dn Mut and Ptn KO mice provide a rationale to investigate if targeting overexpression of Ptn to astrocytes in Ts65Dn Mut mice can rescue or ameliorate the neuronal deficits. We used a viral-mediated strategy to overexpress Ptn, or the control spaghetti monster fluorescent protein (smFP; non-fluorescent modified GFP)^32^, under the minimal GfaABC1d (Gfap) promoter to drive expression in astrocytes (Figure 5A). We delivered virus at P60, after the dendrite and synapse phenotypes are detected at P30, allowing us to evaluate if Ptn can rescue circuit aberrations rather than just prevent the phenotypes from developing. The PHP.eB serotype of adeno-associated virus (AAV) was delivered retro-orbitally to achieve brain-wide targeting (Figure 5SA-B) and allowed for assessment of complex behaviors that involve multiple brain regions. Quantification of virus penetrance and specificity was performed in the VC. Both the Gfap-Ptn and Gfap-smFP viruses targeted ∼60% of Sox9+ cortical astrocytes (smFP: 58.2 ± 6.9%, Ptn: 62.9 ± 1.9%; Figure 5C-D) and were highly specific for astrocytes, with negligible transduction of NeuN+ neurons (smFP: 0.31 ± 0.03% of NeuN+ cells, Ptn: 0.33 ± 0.07% of NeuN+ cells; Figure 5E-F). Four subject groups of the Ts65Dn x Thy1-YFP mice were utilized: WT mice with smFP control (hereafter referred to as “WT”), WT mice with Ptn overexpression (“WT Ptn”), Ts65Dn Mut mice with smFP control (“Ts65Dn”) and Ts65Dn Mut mice with Ptn overexpression (“Ts65Dn Ptn”). YFP-positive mice were used for dendrite and spine quantifications and YFP-negative mice were used for synaptic puncta quantifications. Behavioral analysis was initiated 7 weeks post-injection and tissue was collected at 8 weeks post-injection (∼P120; Figure 5B).

**Figure 5.**
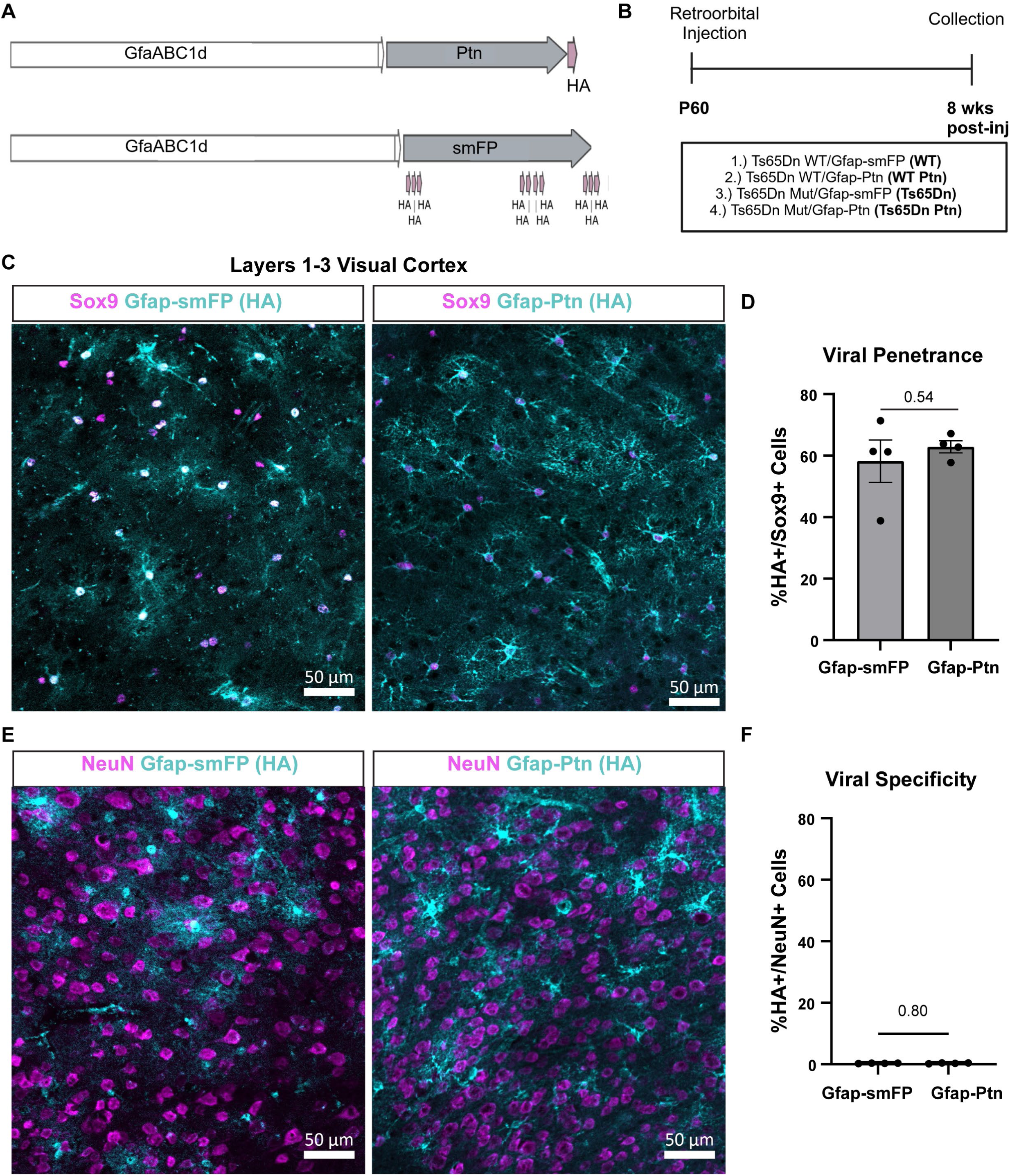
Viral overexpression strategy efficiently and specifically targets astrocytes. (A) Cartoon schematic of targeting viruses. The *GfaABC1d* promoter was used to target astrocytes to either overexpress the control smFP containing 10 HA tags (bottom) or Ptn with an HA tag (top). (B) Timeline of experimental design for viral injection, behavior and collection for histology. (C) Example images of virus labeling with anti-HA (cyan) in Sox9+ (magenta) astrocytes. (D) Quantification of the percentage of Sox9+ astrocytes expressing HA for each virus. (E) Representative images showing that virus (anti-HA, cyan) does not co-label with NeuN (magenta). (F) Quantification of virus expression in neurons shows <0.5% of total neurons have off-target labeling. Scale bars = 50 um. N = 4 mice/group. Statistics by unpaired t-test.

To ask if Ptn overexpression can rescue dendritic phenotypes in Ts65Dn Mut mice we analyzed total dendrite length, dendritic complexity and spine density, in Layer 5 neurons in the VC, as described in Figure 3. Representative filament tracings of dendrites are depicted in (Figure 6A-D). Quantification showed that Ts65Dn dendrites are truncated compared to WT (average of 789.2 ± 51.3 in WT, 591.0 ± 55.5 um in Ts65Dn), reproducing our earlier findings (Figure 3), and overexpression of Ptn fully reversed this deficit (818.3 ± 44.2 um in Ts65Dn Ptn). WT Ptn mice showed a nonsignificant trend towards reduced dendrite length (647.6 ± 21.2 um; Figure 6E) that reached significance when analyzing the combined dendrites from all biological replicates (Figure S6A-B). Sholl analysis revealed that dendritic complexity was also rescued in Ts65Dn Ptn mice, with Ts65Dn dendrites having a decreased area under the curve (average area of 573.4 ± 36.4 in WT, 391.4 ± 42.3 in Ts65Dn), whereas Ts65Dn Ptn dendrites did not differ from WT (area of 552.6 ± 14.9). In this measure dendrites in WT Ptn mice did not significantly differ from WT (area of 479.7 ± 19.7; Figure 6F-G). We next assessed spines, finding that dendrites in Ts65Dn mice had reduced spine density compared to WT (average of 1.66 ± 0.032 spines/um in WT, 1.34 ± 0.091 spines/um in Ts65Dn), as we previously showed (Figure 3), whereas spine density in Ts65Dn Ptn mice was recovered to WT levels (1.69 ± 0.094 spines/um). WT Ptn dendritic spine density was not significantly altered from WT (1.51 ± 0.077; Figure 6H-L; S6C). Investigation of spine morphology showed an increased percentage of filopodia in Ts65Dn mice (average of 9.9 ± 0.8% in WT, 15.1 ± 1.4% in Ts65Dn), which was not impacted by Ptn overexpression (average of 14.2 ± 1.5% in Ts65Dn Ptn). Interestingly, dendrites in WT Ptn mice also had increased filopodia (14.60 ± 0.9%; Figure S6D). Altogether these findings demonstrate that increasing the level of astrocyte-derived Ptn can rescue neuronal dendrite length/complexity and spine density deficits in DS Ts65Dn mice in adulthood.

**Figure 6.**
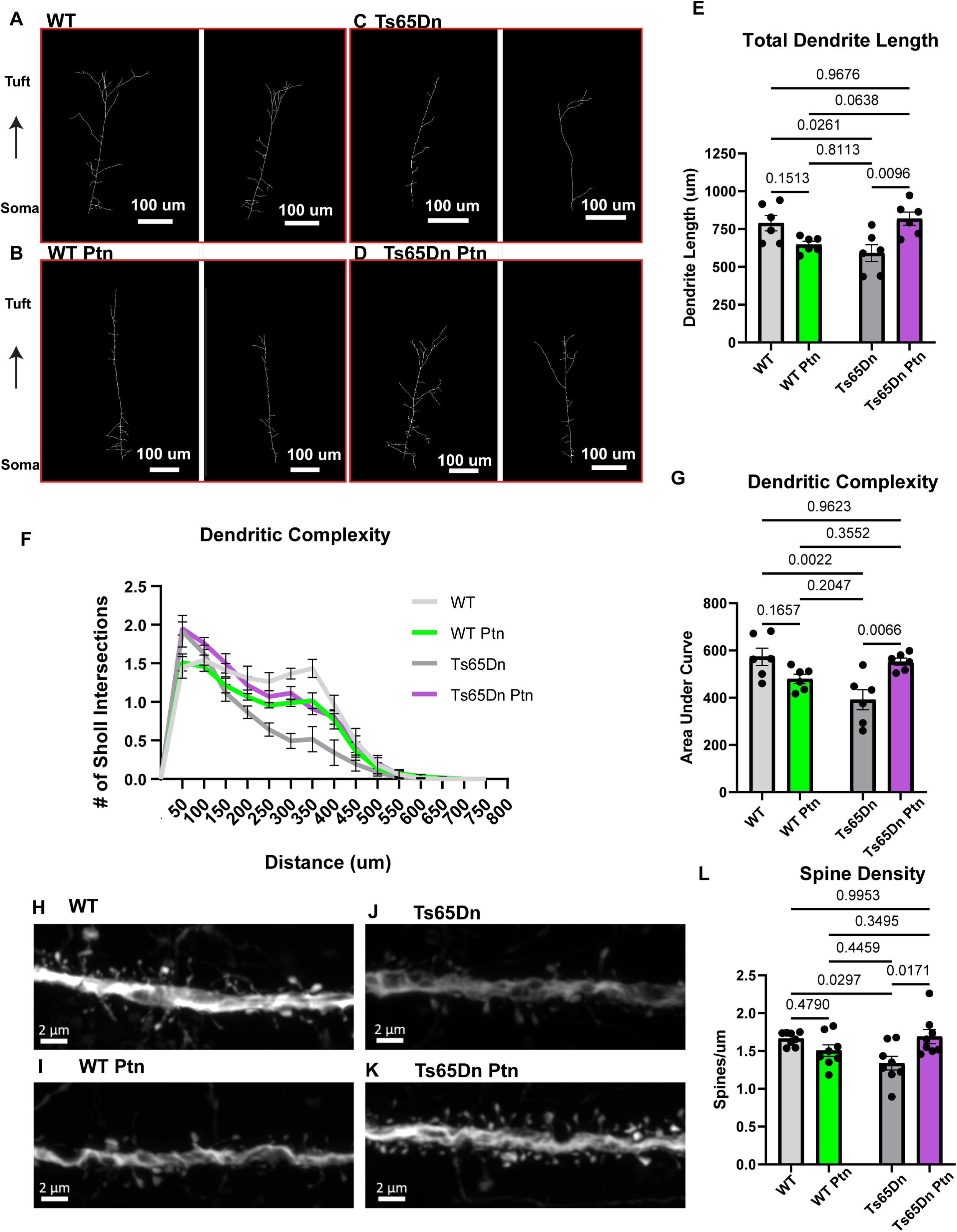
Astrocyte-derived Ptn is a sufficient factor to rescue dendrite and spine phenotypes in Ts65Dn Mut neurons. (A-D). Example filament tracings of dendrites at P120 for each experimental group: WT, WT Ptn, Ts65Dn, Ts65Dn Ptn. The dendrites were traced from the cell body to the dendritic tuft. Scale bars = 100 um. (E.) Total dendrite length is shorter in Ts65Dn compared to WT, and Ts65Dn Ptn dendrites are rescued. N = 6 mice/group. Statistics by two-way ANOVA with Tukey’s test. (F) Sholl curves quantifying dendrite branching points at 50 um increments from the cell body. (G) Area under the curve is decreased in the Ts65Dn group and Ts65Dn Ptn dendrites recover from the reduced branching phenotype. Statistics by two-way ANOVA with Tukey’s. N = 6 mice/group. (H-K) Example images of dendritic spines for WT (H), WT Ptn (I), Ts65Dn (J) and Ts65Dn Ptn (K) groups. Scale bars = 2 um. (L) Spine density is decreased in the Ts65Dn group and Ts65Dn Ptn dendrites recover from the phenotype. N = 8 mice/group. Statistics by two-way ANOVA with Tukey’s.

### Astrocyte-targeted pleiotrophin overexpression improves excitatory synapse phenotypes in DS Ts65Dn mice

To ask if Ptn overexpression can improve the deficits in intracortical synapse number we observed in Ts65Dn mice (Figure 4), we performed immunohistochemical analysis of pre- and postsynaptic markers Vglut1 and GluA2 in WT, WT Ptn, Ts65Dn and Ts65Dn Ptn mice in the VC in Layer 2/3 (Figure 7A-D). This reproduced our earlier finding that Ts65Dn mice have decreased colocalized synaptic puncta (Figure 7A-D) compared to WT (0.66 ± 0.065-fold), while the Ts65Dn Ptn mice showed no significant change compared to WT (0.93 ± 0.092-fold), indicating an improvement of the synaptic phenotype with Ptn overexpression. WT Ptn mice also did not significantly deviate from WT (0.86 ± 0.085-fold; Figure 7G). Vglut1 and GluA2 puncta were quantified individually, showing Ts65Dn mice have a significant decrease in Vglut1 puncta, while Ts65Dn Ptn mice do not significantly deviate from WT (Vglut1: 0.80 ± 0.049-fold in Ts65Dn, 1.01 ± 0.053-fold in Ts65Dn Ptn, 0.96 ± 0.060-fold in WT Ptn; Figure 7E). There are no significant differences in GluA2 puncta in any group (GluA2: 0.84 ± 0.063-fold in Ts65Dn, 1.03 ± 0.076-fold in Ts65Dn Ptn, 0.94 ± 0.060-fold in WT Ptn; Figure 7F). These results demonstrate the ability of Ptn to promote synapse formation in Ts65Dn mice, even at more mature timepoints after developmental synaptogenesis has largely ceased.

**Figure 7.**
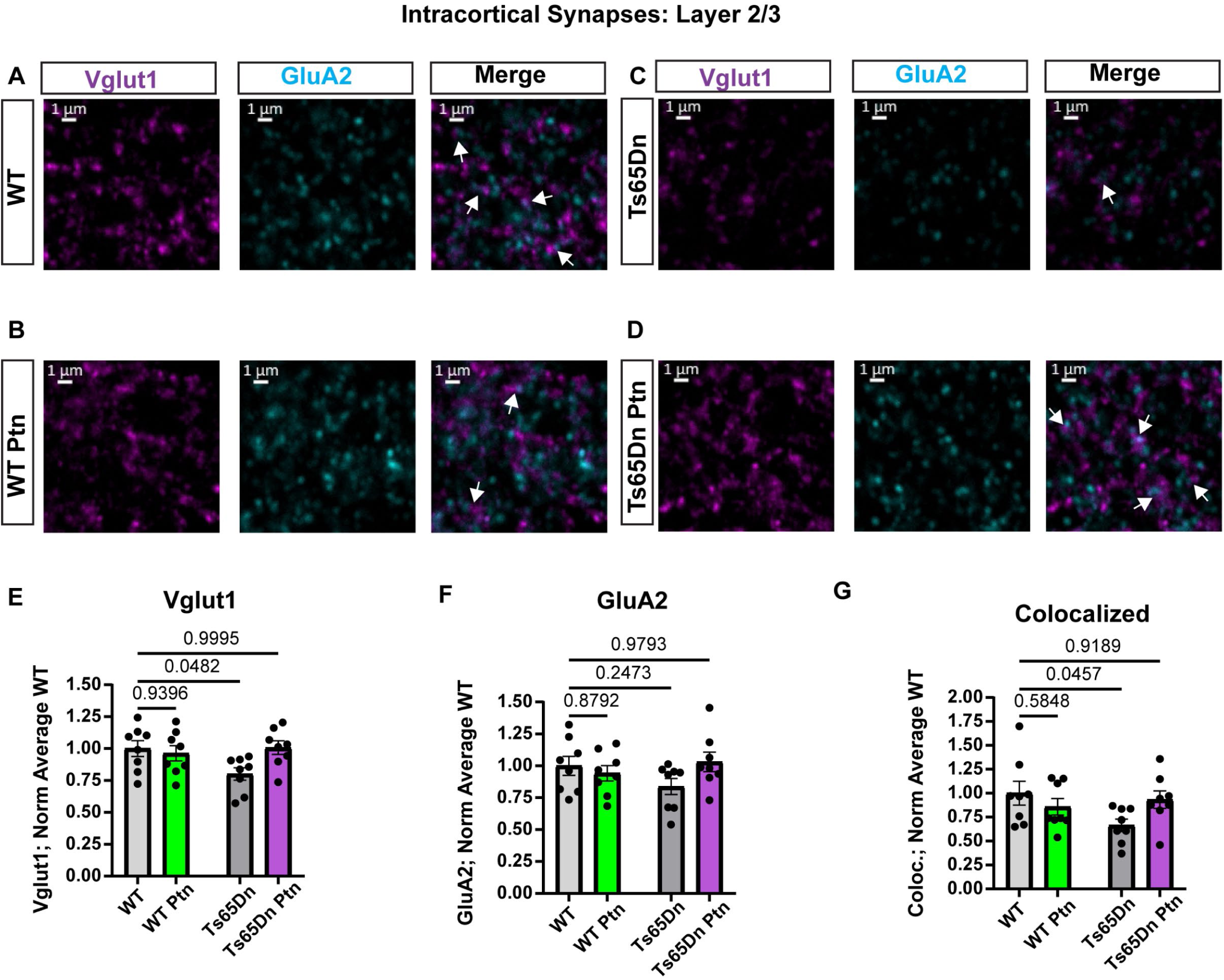
Synaptic phenotypes are improved with Ptn overexpression. (A-D) Example images of Vglut1 (magenta), GluA2 (cyan) and colocalized puncta (white arrows) in Layer 2/3 for WT (A), WT Ptn (B), Ts65Dn (C) and Ts65Dn Ptn (D). Scale bars = 1 um. (E) Vglut1 puncta are decreased in the Ts65Dn group compared to WT. WT Ptn and Ts65Dn Ptn groups have similar Vglut1 levels as WT. (F) GluA2 puncta levels are similar between all groups. (G) Colocalized puncta are decreased in the Ts65Dn group compared to WT. WT Ptn and Ts65Dn Ptn have similar levels of colocalized puncta as compared to WT. All groups normalized to average of WT group. Statistics by two-way ANOVA with Dunnett’s. N = 8 mice/group.

### Astrocyte-targeted pleiotrophin overexpression improves aspects of cognitive function in DS Ts65Dn mice

Our histological evaluation of the effects of Ptn overexpression in astrocytes in Ts65Dn mice demonstrated that Ptn can rescue dendrite morphology and spine density deficits, as well as improve impairments in intracortical excitatory synapse number. To ask if these structural rescues are sufficient to improve performance in behavioral assays, we used a series of behavioral tests where Ts65Dn mice have been reported to show deficits (Figure 8A). Previous work demonstrated that Ts65Dn mice are hyperactive in the open field and have memory impairments in the spontaneous alternation Y maze and fear conditioning assays^33^. Locomotor activity in the open field was assessed over 10 minutes (Figure S7A). We detected a main genotype effect of hyperactivity in Ts65Dn mice in the 2^nd^ 5-minute block, but not the 1^st^ 5-minute block. The observed hyperactivity was not rescued with Ptn (Figure S7B-C). We assessed working memory using the spontaneous alternation Y maze. We found that Ts65Dn mice perform significantly fewer correct alternations than WT (average of 56.9 ± 2.9% in WT, 47.0 ± 1.8% in Ts65Dn) and the Ts65Dn Ptn mice displayed a partial recovery of this phenotype, being not significantly different from WT (51.0 ± 2.2%). WT Ptn mice also performed similarly to WT (57.0 ± 3.0%; Figure 8B). The Ts65Dn mice, regardless of treatment, were hyperactive in this test and entered more arms than WT mice (WT: average of 45.5 ± 2.4 entries, WT Ptn: 49.7 ± 1.9, Ts65Dn: 65.7 ± 4.0, Ts65Dn Ptn: 63.1 ± 3.2; Figure 8C).

**Figure 8.**
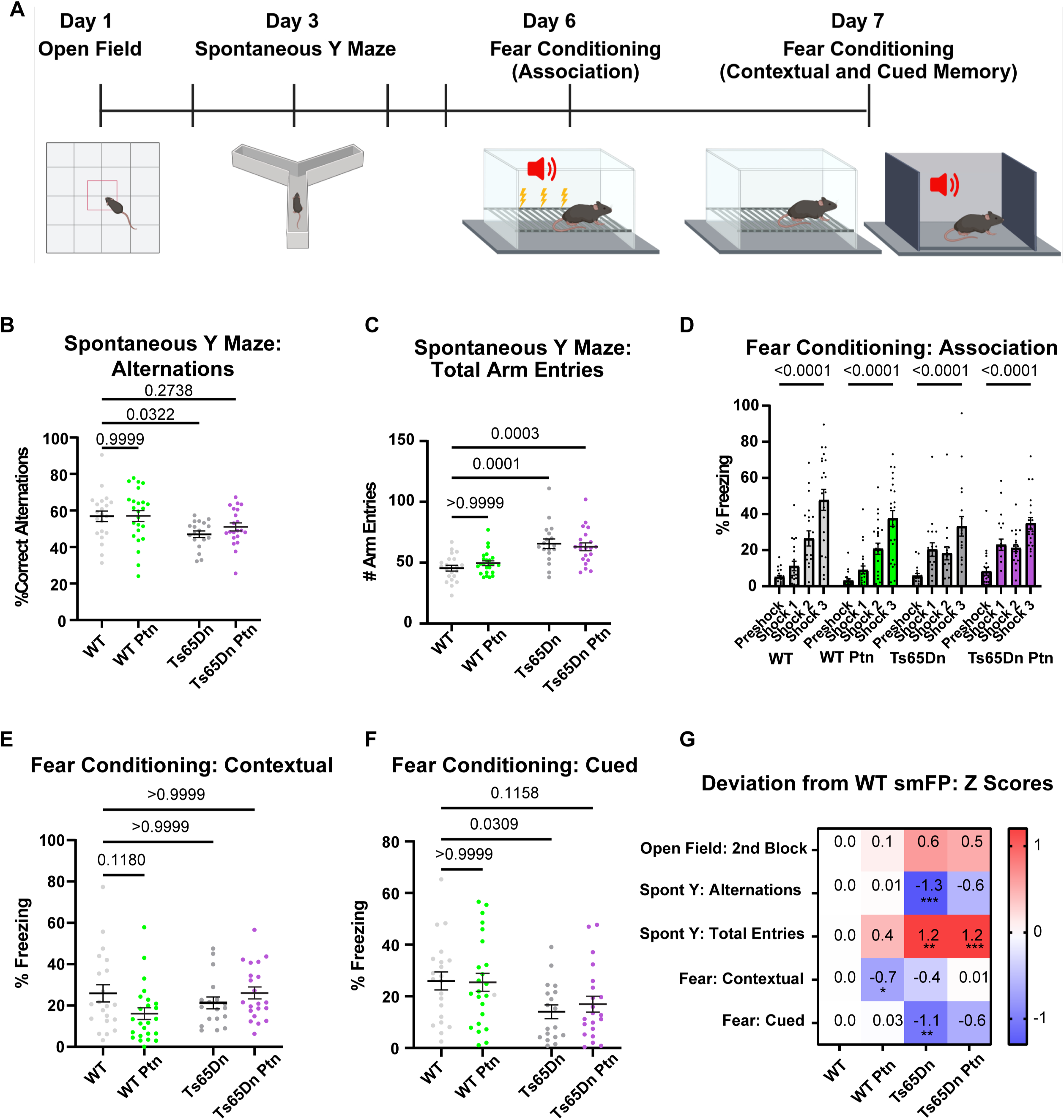
Ptn treatment results in improvement on memory tasks in Ts65Dn Mut mice. (A) Cartoon schematic of behavioral testing schedule. (B) Ts65Dn mice have a decreased percentage of correct alternations in the spontaneous Y maze test. WT Ptn and Ts65Dn Ptn performed at similar levels to WT. Statistics by two-way ANOVA with Dunnett’s. (C) Ts65Dn mice regardless of treatment entered more arms during the test duration than their WT counterparts. Statistics by Kruskal-Wallis with Dunn’s. (D) In fear conditioning mice from each group learned to associate the tone with the foot shock as assessed by the percentage of time freezing after the 3^rd^ shock compared to baseline freezing within a group. Statistics by multiple Mann-Whitney tests with Benjamini, Krieger and Yekutieli correction. (E) We did not observe statistically significant differences between Mut and WT groups in the percentage of time freezing during the contextual memory paradigm. Statistics by Kruskal-Wallis with Dunn’s. (F) Ts65Dn mice exhibited significantly reduced freezing behavior compared to WT in the cued memory paradigm. WT Ptn performed similarly to WT. Ts65Dn Ptn were not significantly different than WT. Statistics by Kruskal-Wallis with Dunn’s. (G) A summary of z-scores for each behavioral test is presented as the deviation of each group from WT. Blue shading signifies a decrease in average z-score and red shading signifies an increase in average z-score. The numeric value of the z-score for each comparison is listed in each box. *** p ≤ 0.001; ** p ≤ 0.01, * p ≤ 0.05. Statistics by two-way ANOVA with Dunnett’s (open field and spontaneous Y maze alternations) or Kruskal-Wallis with Dunn’s (spontaneous Y maze arm entries and contextual and cued fear conditioning). For all tests, For all tests, N = 18-24 mice/group.

We used fear conditioning to test associative learning, as well as contextual and cued memory. We did not observe differences in learning, as each group significantly increased freezing behavior after the 3^rd^ shock compared to baseline (Figure 8D). Twenty-four hours later contextual and cued memory were assessed. For contextual memory, we did not detect any genotype difference in Ts65Dn mice compared to WT (WT: 25.9 ± 4.1%, WT Ptn: 16.0 ±2.8%, Ts65Dn: 21.3 ± 2.8%, Ts65Dn Ptn: 26.1 ± 2.9%; Figure 8E). For cued memory, Ts65Dn mice were significantly impaired compared to WT, showing less freezing behavior (average of 26.0 ± 3.5% in WT, 14.0 ± 2.7% in Ts65Dn), whereas Ts65Dn Ptn mice were not significantly different from WT (17.0 ± 3.1%) and WT Ptn mice performed similarly to WT (25.5 ± 3.5%; Figure 8F). Behavioral measures for each group were also assessed as z-scores deviating from WT, with the three main findings remaining consistent. First, Ts65Dn mice have significantly impaired working memory in the Y maze, but Ts65Dn Ptn mice do not significantly deviate from WT. Second, Ts65Dn mice, regardless of treatment, are hyperactive in the number of arm entries in the Y maze. Third, Ts65Dn mice have significantly impaired cued fear memory while Ts65Dn Ptn are not significantly changed from WT. In addition, when assessed by z-score, WT Ptn mice exhibit significantly reduced freezing in the contextual memory paradigm (Figure 8E). Altogether, these findings demonstrate that Ptn treatment in the DS context can improve function on memory tasks, but does not affect hyperactivity.

## Discussion

We find that astrocyte-derived pleiotrophin is decreased in the Ts65Dn mouse model of DS, and through characterization of Ptn KO mice we identify important neurodevelopmental phenotypes that are mirrored in Ts65Dn Mut mice. We demonstrate that targeting Ptn overexpression specifically to astrocytes in Ts65Dn mice rescues neuronal dendrite length/complexity and spine density phenotypes, as well as partially improves deficits in excitatory synapse number. These histological findings are paired with functional improvements in memory tasks. Importantly, these improvements are seen when delivering Ptn to Ts65Dn mice after phenotypes are already established, showing reversibility. The results of this study highlight the importance of astrocyte regulation of neuronal structure/morphology and neuronal synapses in the context of neurodevelopmental disorders, and provide evidence that manipulating levels of astrocyte-secreted molecules may be an effective therapeutic strategy even after the classical neurodevelopmental windows for synapse formation have closed.

### Roles of astrocyte-derived Ptn in the context of DS and other neurodevelopmental disorders

Stunted dendrite outgrowth and decreased spine density are well-characterized phenotypes in the cortex and hippocampus of DS brains^3–6^. Prior studies on Ptn demonstrated a regulatory role in axon and dendrite outgrowth, and maintenance of spine density in the healthy brain^18,21,34^. However, a link between down-regulated Ptn and DS phenotypes has not been described or functionally investigated and is a major finding of the present study. We identified Ptn as a candidate molecule by mining our recently published dataset describing transcriptomic and proteomic changes to astrocytes from several neurodevelopmental disorders^16^. Ptn protein was found at significantly decreased levels in the secreted fraction from Rett Syndrome and DS astrocytes, with a strong trend towards reduction in Fragile X Syndrome, suggesting that Ptn deficiency may be common to neural circuit impairments observed across multiple neurodevelopmental disorders.

The findings of this study support the concept that astrocyte-secreted proteins can be used as therapeutic tools in DS, and to our knowledge this is the first *in vivo* demonstration of this concept. The present study demonstrates that *in vivo* astrocytes can be manipulated to correct circuit aberrations in a murine model. A prior study found *in vivo* recovery of spine density in the Dp(16) mouse model of DS, with interventions to deplete microglia and reduce inflammation using anti-inflammatory drug treatment. These interventions were performed at P16 and P20, which roughly correlates to childhood to early pubescent years (∼3-15 years) in a human^35,36^. Our intervention overexpressing Ptn in astrocytes at P60 is more equivalent to young adulthood, demonstrating that circuit impairments can be reversed. We chose to use AAV-mediated gene delivery for purposes of potential clinical translation. AAV gene therapies exhibit low toxicity coupled with the ability to drive prolonged gene expression^37^. An AAV gene therapy is currently in Phase 1/2 clinical trials for Rett Syndrome (REVEAL Adult Study, Clinical Trial #NCT05606614) for purposes of normalizing levels of the MECP2 gene that causes the disorder. Importantly, our findings highlight the therapeutic potential of using AAV-mediated delivery of astrocyte-secreted proteins as a tool to correct deficits in circuit connectivity and function in neurodevelopmental disorders that are not caused by a single gene mutation.

### Roles of Ptn in neurotypical development

In the present study we found remarkable overlap in the neurodevelopmental phenotypes observed in Ts65Dn Mut and Ptn KO mice, but there were also notable differences. Both mouse models had decreased levels of intracortical excitatory synapses in the upper cortical layers. However, Ptn KO mice additionally showed reduced levels of Vglut2 in Layer 1 which correspond to the thalamocortical axon projections^38^. Given the demonstrated roles of Ptn in neurite outgrowth, we hypothesize that Ptn may act as a guidance factor for the long-range thalamocortical projections during early cortical development. These Vglut2 deficits were present at P30 and P120, implying that this is not merely a developmental delay, and rather could be related to impaired navigation to the target innervation zone during development. It will be interesting in future studies to investigate if Ptn KO mice have any related functional deficits, for example in visual acuity.

Secondly, the total dendrite length and complexity phenotypes in Ts65Dn mice are present at both P30 and P120, but Ptn KO mice recover from this phenotype by P120. These findings suggest that in the case of genetic loss of *Ptn*, the phenotype may be related to a developmental delay, or there may be other local compensatory factors that are able to recover dendrite outgrowth. For example, midkine is a closely related paralogue of Ptn that has overlapping, but not identical, functions^39^. A genetic loss of *Ptn* could be compensated for by midkine in the Ptn KO model, whereas an incomplete loss of Ptn in Ts65Dn Mut mice may not trigger the same compensatory mechanisms.

### Importance of maintaining homeostatic levels of astrocyte-derived factors

Many astrocyte-derived factors have been shown to promote synapse formation and maturation. However, most of these studies were performed with KO mouse models showing that loss of the factor of interest is detrimental to circuit function^10,31,40–42^. There is currently a knowledge gap regarding circuit outcomes *in vivo* when astrocyte-derived factors are expressed at levels higher than what is physiological, rather than to rescue a deficit. In this study we find that overexpression of Ptn in the WT context impairs total dendrite length. We also observed decreased freezing in Ptn-treated WT mice in the contextual memory assay. Work from the hippocampus has shown that Ptn is an injury-induced factor. *Ptn* mRNA is decreased transiently following forebrain ischemic insult or excitotoxic injury from kainic acid injection, but four days post-injury *Ptn* is up-regulated^43^. The transient decrease in *Ptn* expression after excitotoxic injury suggests it may be down-regulated in response to high levels of activity but up-regulated when tissue repair begins and formation of new synapses is prioritized. We hypothesize that in a non-diseased context, up-regulation of Ptn may have harmful effects on synaptic transmission, but in a diseased context where synapses are not maintained at homeostatic levels then Ptn overexpression may be beneficial, as we have demonstrated in the context of DS. These are important considerations when evaluating astrocyte-secreted proteins as potential therapeutics during safety studies in clinical trials that assess healthy subjects.

In conclusion we present evidence that Ptn is an astrocyte-secreted protein that promotes development of neuronal dendrite morphology and synaptic circuitry during neurotypical development. We demonstrate that Ptn is down-regulated in astrocytes from a commonly used mouse model of DS, and use viral-mediated overexpression to fully rescue dendrite and spine deficits, partially restore excitatory intracortical synapse alterations, and improve behavioral outcomes. This work opens investigative avenues for astrocyte-based therapeutics that may be relevant to multiple neurodevelopmental disorders.

## Supporting information

Supplemental Figures with legends

Supplemental statistics file for Figure 1 and Figure S1

Associated statistics file for Figure 2 and Figure S2

Associated statistics file for Figure 3 and Figure S3

Associated statistics file for Figure 4 and Figure S4

Associated statistics file for Figure 5

Associated statistics file for Figure 6 and Figure S6

Associated statistics file for Figure 7

Associated statistics file for Figure 8 and Figure S8

## Supplemental Information

Supplemental information includes 7 figures and 8 supplemental statistics tables.

## Acknowledgements

We thank Cari Dowling and Joseph Hash for technical assistance with the mouse colony and T.F. Vogt for sharing the Ptn KO mice. We thank Krissy Lyon for cloning of the GfaABC1d-smFP plasmid. Biorender was used to generate cartoon schematics. This work is supported by the Chan Zuckerberg Initiative (NJA) and NIH NINDS F32NS117776 (ANB). MM was supported by STARTneuro program at UCSD (NIH BP-ENDURE grant R25NS119707). This work was supported by Core Facilities of the Salk Institute (In Vivo Scientific Services (Animal Resources Department); Gene Transfer, Targeting and Therapeutics Viral Vector (GT3) Core: supported by NIH NCI, NEI and NINDS; Advanced Biophotonics Core: supported by NIH NCI CCSG: P30 014195 and the Waitt Foundation).

## Author Contributions

ANB and NJA conceived the project, designed experiments, analyzed data, and wrote the manuscript. ANB, QNA and MM performed experiments and analyzed data.

## Declaration of interests

The authors declare no competing interests.

## STAR Methods

### Resource Availability

#### Lead contact

Requests for additional information and sharing of resources and reagents should be directed to the lead contact, Nicola J. Allen.

#### Materials availability

The GfaABC1d-smFP-HA and GfaABC1d-Ptn-HA AAV PHP.eB viral constructs were generated for this study. Original plasmids are available upon request.

### Experimental model and subject details

#### Mice

All animal work was approved by the Institutional Animal Care and Use Committee (IACUC) of the Salk Institute for Biological Studies. Mice were housed in the Salk Institute Animal Resources Department on a light cycle of 12 hours light:12 hours dark with access to food and water *ad libitum*. Mice of both sexes were used for all experiments between the ages of postnatal day (P)0 and P120. For experiments using developmental timepoints, littermates were used. For P120 experiments within-colony controls were used.

### Aldh1L1-EGFP mice

Tg(Aldh1l1-EGFP)OFC789Gsat/Mmucd (Stock 011015-UCD) were used for analysis of developmental *Ptn* mRNA expression in neurotypical development (in situ data).

### Ptn KO mice

Ptn WT and KO mice were used for *in vitro* neuron and astrocyte cell culture experiments. Homozygous WT and homozygous KO crosses were used to obtain larger numbers of pups of each genotype for *in vitro* assays. For propagation and maintenance of the Ptn line, heterozygous crosses were used. WT and KO progeny from the heterozygous crosses were used for the synaptic characterization experiments. Mice were a kind gift from T.F. Vogt at Princeton University and were originally described in^27^.

### Ptn x Thy1-YFP mice

The Ptn line was crossed to the B6.Cg-Tg(Thy1-YFP)HJrs/J line (Jax Stock 003782) to sparsely label Layer 5 pyramidal neurons^29^. The line was maintained as heterozygous for YFP. The YFP+ progeny that were WT for Ptn or KO for Ptn were used for characterization of dendrite length, dendritic complexity and spine density.

### Ts65Dn mice

B6EiC3Sn.BLiA-Ts(1716)65Dn/DnJ mice were originally described in^44^ and obtained from Jax (Stock 005252). These mice were used for synaptic characterizations. The line was propagated and maintained by crossing female mutants to wildtype males on the same background due to male sterility.

### Ts65Dn x Thy1-YFP mice

The Ts65Dn line was crossed to the B6.Cg-Tg(Thy1-YFP)HJrs/J line (Jax Stock 003782) to obtain YFP-labeled Layer 5 pyramidal neurons. The line was maintained as heterozygous for YFP. Due to the Thy1-YFP mice being on a different genetic background than the Ts65Dn mice, only the F1 progeny were used for experiments to avoid extensive backcrossing. Thy1-YFP breeders were imported from Jax as needed. The euploid and trisomic YFP-positive progeny were used for characterization of dendrite length, dendritic complexity and spine density experiments, as well as the associated rescue experiments. The euploid and trisomic YFP-negative progeny were used for the synaptic characterization and rescue experiments.

#### Cell cultures

Primary astrocyte and neuron cultures were obtained using the Miltenyi MACS isolation systems for astrocytes and neurons described in detail below. P6-7 mouse pups (4 pups/experiment) were used for primary astrocyte isolation. P0-1 mouse pups (2 pups/experiment) were used for primary neuron isolation.

### Primary astrocyte cultures

The cortices were dissected and meninges removed in ice-cold DPBS. The cortices were cut into ∼1mm^3^ pieces using a razor blade to allow for optimal enzymatic digestion. Papain digestion was performed according to the kit instructions (Miltenyi MACS Neural Tissue Dissociation Kit 130-092-628). Digestion was performed using the gentleMACS Octo Dissociator with Heaters (Miltenyi 130-096-427) using the 37C_NTDK_1 protocol provided by Miltenyi with a total time of 22 minutes. Cells were centrifuged at 4C 300 g for 10 minutes and then resuspended in AstroMACS buffer and passed through a 70-um basket filter to remove clumps. Astrocytes were subsequently isolated using the Miltenyi MACS Anti-ACSA-2 Microbead kit (Miltenyi 130-097-678) according to the manufacturer’s instructions, with slight modifications to deplete myelin debris and microglial cells prior to positive astrocyte selection. Myelin was depleted using a 10-minute incubation at 4C with anti-myelin beads (Miltenyi 130-104-257). The MS magnetic separator column was then used to deplete the myelin from the cell suspension via magnetic separation. The flow-through was collected and centrifuged at 4C 300 g to remove excess beads. Microglia depletion was performed using anti-CD11b beads (Miltenyi 130-093-634) for an incubation period of 10 minutes at 4C. The MS magnetic separator column removed the microglia from the cell suspension via magnetic separation. The flow-through was collected and centrifuged again at 4C 300 g to remove excess beads. The supernatant was collected and subsequent positive selection for astrocytes was performed. First a 10-minute incubation with FcR blocking beads was used to minimize any nonspecific binding. ACSA2-conjugated beads were then used to positively select for astrocytes during a 15-minute incubation period at 4C (FcR and ACSA2 beads in kit, Miltenyi 130-097-678). The MS column was used to magnetize the ACSA2-beads and collect the astrocytes. The magnetized astrocytes were plunged into a new tube with AstroMACS buffer and centrifuged again at 4C 300 g to remove excess beads. Cells were counted using Trypan Blue solution (Gibco 15250-061) and plated in 6-well plates (Falcon 353046) coated with poly-D-lysine (Sigma P64047) at a density of 40,000-60,000 cells per well. Astrocytes were grown to confluency over 7 days *in vitro* (DIV) in astrocyte growth medium containing: 50% DMEM (Thermo Fisher Scientific 11960044), 50% Neurobasal (Thermo Fisher Scientific 21103049), Penicillin-Streptomycin (LifeTech 15140-122), Glutamax (LifeTech 35050-061), sodium pyruvate (LifeTech 11360-070), N-acetyl-L-cysteine (Sigma A8199), SATO (containing transferrin (Sigma T-1147), BSA (Sigma A-4161), progesterone (Sigma P6149), putrescine (Sigma P5780) and sodium selenite (Sigma S9133)), and heparin binding EGF like growth factor (HbEGF,5 ng/mL; R&D systems 259-HE/CF). Astrocyte cultures were maintained in a humidified incubator at 37C/10% CO2.

### Collection of astrocyte conditioned media (ACM)

The growth media was removed from the 6-well plates after 7DIV and the wells were washed three times with warm Dulbecco’s Phosphate Buffered Saline (DPBS; HyClone SH30264.01). Astrocytes were then cultured in minimal low protein conditioning media containing: 50% DMEM, 50% Neurobasal, penicillin-streptomycin, glutamax, sodium pyruvate, and HbEGF. The culture was maintained in a humidified incubator at 37C/10% CO2 for 4DIV. The conditioned media was concentrated using Vivaspin-6 centrifugal concentrators with a molecular weight cut-off of 10 kDa (Sartorius VS0602). Protein concentration in the ACM was assessed using a Qubit 4 fluorometer (Invitrogen Q33238) and the Qubit protein assay kit (Invitrogen Q33211). The ACM for neuronal dendrite outgrowth assays was stored at 4C for no longer than 1 week before use. The ACM used for the neuronal dendrite outgrowth assays was produced from either WT astrocytes or Ptn KO astrocytes.

### Primary neuron cultures

The cortices were dissected and meninges removed in ice-cold DPBS as described above. Papain digestion was performed the same as for primary astrocyte isolation. Cells were centrifuged at 4C 300 g for 10 minutes and then resuspended in cold DPBS and passed through a 70-um basket filter to remove clumps. Neurons were subsequently isolated by biotin-conjugated magnetic bead depletion of non-neuronal cells using the Miltenyi Neuron Isolation Kit according to the manufacturer instructions (Miltenyi 130-115-389). Depletion with the non-neuronal antibody cocktail was performed for 5 minutes at 4C followed by centrifugation at 300 g (4C) for 10 minutes. Anti-biotin beads were then used to remove the non-neuronal cells during a 10-minute incubation at 4C. The cell suspension was applied to the Miltenyi LS column and the flow-through (now a neuron-enriched cell suspension) was collected. Trypan Blue was used to count cells and the neurons were plated on glass coverslips (12 mm diameter, Carolina Biological Supply 633029) coated with poly-D-lysine and laminin (R&D 3400-010-01) in 24-well plates (Falcon 353047) at a density of 40,000-60,000 cells per well. Neurons were cultured in either minimal media (MM), MM + WT ACM, MM + Ptn KO ACM or Ptn KO ACM + 0.25 ug recombinant Ptn (“Rescue ACM”; see below). The MM contained 50% DMEM, 50% Neurobasal, Penicillin-Streptomycin, Glutamax, sodium pyruvate, N-acetyl-L-cysteine, insulin (Sigma I6634), triiodo-thyronine (Sigma T6397), SATO, B27 (containing L-carnitine hydrochloride (Sigma C0283), ethanolamine (Sigma E9508), D(+)-galactose (Sigma G0625), putrescine (Sigma P5780), sodium selenite (Sigma S9133), corticosterone (Sigma C2505), linoleic acid (Sigma L1012), lipoic acid (Sigma T1395), progesterone (Sigma P8783), retinyl acetate (Sigma R7882), retinol (Sigma 95144), D,L-alpha tocopherol acetate (Sigma T3001), bovine albumin (Sigma A4161), catalase (Sigma C40), glutathione (Sigma G6013), insulin (Sigma I6634), superoxide dismutase (Sigma S5395), transferrin (Sigma T5391), triiodol-l-thyronine (Sigma T6397)), and forskolin (Sigma F6886). Cortical neuron cultures were maintained in a humidified incubator at 37C/10% CO2.

#### Method Details

##### Treatment of neuronal cultures for dendrite outgrowth assays

Primary neurons isolated from WT mice were cultured for 4DIV in either MM, WT ACM, Ptn KO ACM or Ptn KO ACM + 0.025 ug recombinant human Ptn (Rescue ACM, R&D Systems 252-LP-050). 1.5 ug/well of ACM was added to MM for the conditions receiving ACM treatment. The neurons were treated twice with ACM, at 0DIV and again at 2DIV. At 4DIV the neurons were collected for analysis (see methods details for immunocytochemistry below).

##### Dendrite outgrowth assay/immunocytochemistry

The cover slips with neurons were washed three times with warm DPBS followed by 20 minutes fixation with room temperature 4% paraformaldehyde (PFA, Electron Microscopy Sciences 50980487) diluted in PBS. Coverslips were washed three times with room temperature DPBS and subsequently blocked/permeabilized in a solution of 50% goat serum (Life Technologies 16210072)/50% antibody buffer (containing 150 mM sodium chloride (Sigma S5886); 50 mM Trizma base (Sigma T6066), 100 mM L-lysine (Sigma L5501), 1% bovine serum albumin (Sigma A2153), pH 7.4) and 0.5% Triton X-100 (Sigma T9284). The blocking step was performed for 30 minutes. Coverslips were then washed once in PBS and incubated overnight at 4C with antibodies against Map2 (1:2500, chicken, EnCor Biotechnologies CPCA-Map2) and Ctip2 (1:500, rat, Abcam ab18465). The next day coverslips were washed three times in PBS and incubated for two hours with appropriate secondary antibodies (goat anti-chicken 488, 1:500, Thermo Fisher Scientific A-11039) and goat anti-rat 555, 1:500, Thermo Fisher Scientific A-21434) in a solution of antibody buffer with 10% goat serum. Coverslips were washed three times in room temperature PBS and then stained with 4′,6-diamidino-2-phenylindole (DAPI, 1:1000, Thermo Fisher Scientific 62247) for 5 minutes. Excess DAPI was washed off with one PBS wash and then coverslips were mounted onto glass slides using Fluoromount G (Southern Biotech 0100-01). Imaging was performed with a 20x objective on a Zeiss LSM 700 confocal microscope. Z-stacks were collected with 1.0 um steps and a total of 5 steps. The maximum intensity projection images were used to assess total dendrite length and complexity. For each condition a total of 3 coverslips (technical replicates) were imaged per experiment. Ten images were acquired for each coverslip for a total of 30 images per condition in each experiment. Each Ctip2+ cell in the image was quantified if its entirety of dendrite labeling was captured in the image. Quantification of total dendrite length and Sholl analysis for dendritic complexity was performed in Imaris v9.2.0 (Oxford Instruments) using the filament tracing tool. Sholl analysis was performed with a resolution of 10 um. Four independent experiments were performed and analysis of dendrite length was conducted on both the average of cells within an experiment (Figure 2) and on a per cell basis (Figure S2) with the same results.

##### Western blotting

Western blotting was used to determine the physiological range of Ptn secretion from astrocytes. For determination of the physiological range of Ptn secretion from cultured astrocytes, four independent culture experiments were used. The ACM total protein concentration was assessed using the Qubit Protein Assay kit and ACM was loaded into the wells at inputs of 3 ug, 6 ug and 9 ug. Recombinant human Ptn protein was added into the wells at: 0.1 ug, 0.2 ug and 0.3 ug. The ACM and recombinant protein samples were heated in reducing loading dye (Thermo 39000) at 55C for 45 minutes. The samples were separated on 4-12% Bis-Tris gradient gels (Invitrogen NW04122BOX) at 80V for 1.5 hours in MOPS SDS buffer (Invitrogen B0001.02). Proteins were transferred to PVDF membrane (Millipore IPFL00010) at 100V for 1 hour in an ice bucket using Tris-Glycine buffer (Thermo Fisher Scientific 28363) with 20% methanol (Fisher A412-4). The membrane was blocked in blocking buffer containing 0.1% Casein (BioRad 1610782) in Tris Buffered Saline (TBS, BioRad 105300272) for one hour at room temperature on a shaker. The membrane was incubated overnight at 4C in blocking buffer with anti-Ptn antibody (1:500, goat, R&D Systems AF-252-PB). The next day the membrane was washed three times for ten minutes each in TBS with 0.1% Tween-20 (Sigma P9416) followed by secondary antibody (diluted 1:20,000 in blocking buffer + 0.1% Tween, donkey anti-goat 680, Invitrogen A21084) incubation for one hours at room temperature on a shaker. The membrane was washed three times for ten minutes each in TBS and the bands were visualized using the Odyssey Clx Infrared Imager (Li-Cor) and the band intensity was analyzed using Image Studio software (Li-Cor). The band intensities for WT ACM inputs were compared as ratios to the known input of recombinant Ptn to get an estimate of the fraction of Ptn protein of total protein in the ACM. These experiments yielded a range of 0.01-0.048 ug of Ptn in 1.5 ug of WT ACM. Based on this range, the final value of 0.025 ug Ptn was chosen for the dendrite outgrowth experiments described above. Molecular weights were visualized using Precision Plus All Blue Protein standard loading dye (BioRad 1610373.

##### Cloning of plasmids for viral vectors

Untagged mouse Ptn cDNA clone in a PCMV6-Kan/Neo vector was obtained from oriGene (Accession No. NM_008973, Origene MC204839). The pCAG_smFP-HA vector was obtained from Addgene (#59759). Ptn and smFP-HA were separately inserted into pZac2.1 GfaABC1d-tdTomato vector obtained from Addgene (#44332) after removing tdTomato sequence. The In-Fusion HD cloning kit (Takara 638909) was used according to the manufacturer’s instructions. The GfaABC1d-tdTom vector was linearized at the NheI and XbaI (New England Biolabs R3131S, R0145S) restriction sites for three hours at 37C and the linearized vector was purified from an agarose gel and cleaned using Nucleospin columns (Macherey-Nagel 740609.50). The Ptn cDNA was PCR amplified and an HA tag added to the C-terminus using the following primer sequences with 15-16 bp overlapping sequences to the linearized vector:

Forward (5’ to 3’): ctcactataggctagcatgtcgtcccagcaatatca

Reverse (5’ to 3’): caggagaagatgctggattatccgtatgatgttccggattatgcatagtctagagtcga

The PCR product was subsequently gel purified and cleaned with the Nucleospin columns. The In-Fusion HD cloning enzyme premix was used to insert the Ptn-HA sequence into the vector.

The smFP-HA was cloned into the GfaABC1d-tdTom vector using the same procedure. The primer sequences used for smFP are as follows:

Forward (5’ to 3’): ctcactataggctagcatgtacccttatgatgtgcccgattatgct

Reverse (5’ to 3’): gtacgatgtcccggactacgcttaatctagagtcgacccgg

Stellar competent E. Coli cells (Takara 636763) were then transformed with the plasmids via heat-shock according to the manufacturer’s instructions. Bacteria were screened using carbenicillin resistance (100 ug/mL; Teknova C2130). The colonies were expanded in an overnight LB culture. The DNA was purified using an endotoxin-free plasmid MaxiPrep kit according to the manufacturer’s instructions (Qiagen 12362).

##### Viral vector generation

Plasmids were sequenced for accuracy using Eton Sequencing before packaging into AAV at the Salk Institute Viral Vector (GT3) core facility. Viruses were packaged as AAV PHP.eB serotype to allow for systemic retroorbital delivery^45^ in order to obtain brain-wide transduction of astrocytes.

##### Retroorbital virus injections

Mice at P60 were anesthetized with 2-3% isofluorane in oxygen by constant flow through a nose cone. Virus was injected via retroorbital administration at a dose of 1.0E12 viral genome copies in a total injection volume of 160 ul. The virus was diluted in DPBS just prior to injection. Mice were allowed to recover from anesthesia and then returned to the housing room.

##### Tissue collection and preparation for histological experiments

Mice were deeply anesthetized with intraperitoneal injection of a mixture of 100 mg/kg Ketamine (Victor Medical Company 1699053) and 20 mg/kg Xylazine (Victor Medical Company 1264078) and transcardially perfused. For fixed tissue collection, mice were perfused first with room temperature PBS followed by 4% PFA. The brains were rapidly dissected and post-fixed overnight in 4% PFA (Electron Microscopy Sciences 19208) at 4C. After 24 hours the brains were moved to 30% sucrose (Invitrogen 1503-022) for 24-48 hours. The brains were then embedded in TFM (Electron Microscopy Sciences 72593) and frozen in a dry ice/ethanol slurry and stored at −80C until use. For fresh frozen tissue collection, mice were perfused with room temperature PBS and the brain was rapidly dissected and embedded in OCT (TissueTek 4583). The brains were frozen in a dry ice/ethanol slurry and stored at −80C until use as described above.

##### Single molecule fluorescent in situ hybridization (smFISH)

smFISH was used to characterize the developmental pattern of *Ptn* expression in astrocytes, confirm loss of *Ptn* in Ptn KO mice and validate the reduction of Ptn mRNA in Ts65Dn astrocytes. FISH was performed with the RNAScope Fluorescent Multiplex Detection kit (ACD 320851) according to the manufacturer’s instructions for fixed and fresh frozen experiments (protocols ACD 320535-TN and ACD 320513-USM, respectively).

#### Characterization of developmental Ptn expression in astrocytes

Sagittal sections of brain were sliced at 16 um using a cryostat (Hacker Industries OTF5000). Sections were obtained from Aldh1L1-EGFP mice at P7, P14 and P30. An epitope retrieval step was performed at 95C for 20 minutes using the ACD epitope retrieval buffer. Slides were then washed once in PBS and dehydrated in 100% ethanol before beginning protease treatment. Protease III was used for P7 samples and Protease IV was used for P14 and P30 samples. All protease treatment was performed for 30 minutes at 40C in the hybridization oven. *Mus*

*musculus Ptn* probe (ACD 486381) was hybridized for 2 hours at 40C. Amplification steps were performed according to the manufacturer’s instructions without deviation. *Ptn* probe was conjugated to the 550-channel. After completion of the probe labeling, immunohistochemical labeling was performed to amplify the transgenic GFP signal in astrocytes. Briefly, slides were blocked and permeabilized in blocking buffer containing 10% goat serum in PBS with 0.2% Triton for 1 hour at room temperature. Slides were then incubated overnight at 4C with anti-GFP antibody (1:500, chicken, Novus Biological NB100-1614) diluted in an antibody dilution buffer containing 5% goat serum in PBS with 0.2% Triton. The next day slides were washed in PBS three times for 5, 10 and 15 minutes. Secondary antibody was applied (1:500, goat anti-chicken 488) for 2 hours at room temperature in antibody dilution buffer. Slides were washed three times for five, ten and fifteen minutes. DAPI was applied (1:1000) for five minutes. Slides were washed once in PBS and then coverslips (24 x 60 mm and 1.5 thickness, Fisher 12541037) were mounted with Fluoromount G. Five independent experiments were performed. Imaging was performed using a 40x objective on a Zeiss LSM 700 confocal microscope. Laser and gain intensities were maintained as constant for each image. Tiled z-stacks were obtained from the visual cortex with 0.4 um z-steps and a total of 12 steps, focusing on the upper layers due to description of *Ptn* as an upper layer astrocyte marker ^26^. Three slices were imaged for each mouse. Layer 1 and Layer 2/3 were segmented out as separate images in FIJI using the DAPI channel. Images were quantified using a custom macro in FIJI that masks the astrocyte cell bodies and measures the thresholded mRNA signal in each cell as previously described ^42^. The threshold was maintained as constant for each image within an experiment. Example images in Figure 1 show the maximum intensity projection for each timepoint. Overview images in Figure S1 were collected using a 20x objective. A z-step size of 3.5 um was used with a total of 9 steps. Tiled images of the entire cortex were collected and the image shown is a maximum intensity projection.

#### Confirmation of Ptn KO

Coronal sections of 16 um were obtained from WT and *Ptn* KO mice at P30. Epitope retrieval, FISH and immunohistochemical labeling for S100b was performed on fixed tissue in the same manner as described above. Images were obtained on the LSM 700 confocal microscope using a 40x objective. Tiled z-stacks were collected from the visual cortex with 1 um z-steps and a total of 7 steps. Constant laser and gain settings were used for WT and KO pairs. Images shown are maximum intensity projections.

#### Validation of reduced *Ptn* expression in Ts65Dn Mut astrocytes

smFISH was used to validate that *Ptn* expression is reduced *in vivo* in Ts65Dn Mut astrocytes. Sagittal sections of brain from Ts65Dn WT and Mut littermates at P14 were sliced at 16 um. Dual-probe smFISH was performed on fresh frozen tissue for these experiments, as recommended by the manufacturer. Slides were fixed in 4% PFA at 4C for 15 minutes and then sequentially dehydrated in 50, 70 and 100% ethanol. Protease IV treatment was applied for 8 minutes at room temperature. Hybridization was performed as described above with *Ptn* probe and *Mus musculus Slc1a3* probe (ACD 430781-C2) to label astrocytes. Amplification steps were performed as described above and not deviating from the manufacturer’s instructions. *Ptn* was conjugated to the 550-channel and *Slc1a3* was conjugated to the 488-channel. Five independent experiments were performed. Imaging was performed using a 20x objective on a Zeiss LSM 700 confocal microscope. Laser and gain intensities were maintained as constant for each image. Three slices were imaged for each mouse. Tiled z-stacks were obtained from the visual cortex with 0.4 um z-steps and a total of 12 steps, again focusing on the upper layers. Layer 1 and Layer 2/3 were segmented in FIJI as described above. The same custom macro was utilized for quantification as described above, but masking on *Slc1a3*+ signal. Example images in Figure 1 show the maximum intensity projection. The fresh frozen protocol yielded more variability between independent experiments so the *Ptn* signal in Ts65Dn Mut astrocytes was normalized to the WT values within each experiment.

### Validation of viral constructs

The penetrance and specificity of the GfaABC1d-Ptn-HA and GfaABC1d-smFP-HA viruses were quantified. Viruses were injected and expressed for two months before collection of the brains. Coronal sections of 16 um were obtained from mice injected with each virus. TFM was removed with a PBS wash. Slices were blocked and permeabilized in blocking buffer (10% goat serum in PBS with 0.2% Triton) for one hour at room temperature. Overnight staining with antibody dilution solution (5% goat serum in PBS with 0.2% Triton) was performed at 4C. Antibodies for the astrocyte marker Sox9 (1:1000, rabbit, Abcam ab185966) and the neuronal marker NeuN (1:500, rabbit pre-conjugated to 647, Abcam ab190565) were used in separate tissue sections and each co-labeled with anti-HA antibody (1:500, rat, Sigma 11867423001). The next day slices were washed three times in PBS with 0.2% Triton for 5, 10 and 15 minutes. Secondary antibody labeling was performed in 5% goat serum in PBS with 0.2% Triton for two hours at room temperature. Appropriate secondary antibodies were used (1:500, goat anti-rabbit 488, Thermo Fisher Scientific A11034; 1:500, goat anti-rat 555, Thermo Fisher Scientific A21434). Washes in PBS with 0.2% Triton were performed three times for 5, 10 and 15 minutes. DAPI was applied (1:1000) for 5 minutes followed by a PBS wash. Coverslips were mounted with Fluoromount G. Quantification of viral penetrance and specificity was performed in four mice for each virus. Images were obtained on an LSM 700 confocal microscope using a 20x objective. Tiled z-stacks in the visual cortex were acquired with 2.0 um z-step increments and a total of 8 z-steps. Three slices/mouse were imaged. The images were stitched and maximum intensity projections were created in Zen software. The images were then imported into FIJI for cell counting analysis using the cell counter plug-in. The number of Sox9+ and NeuN+ cells were counted in each image to obtain the total number of astrocytes and neurons, respectively. Then the number of HA+/Sox9+ and HA+/NeuN+ cells were counted. The viral penetrance was quantified as the percentage of Sox9+ cells that were also HA+ for each virus. The off-target expression in neurons was quantified as the percentage of NeuN+ cells that were also HA+. The hippocampus and lateral amygdala were also imaged to show viral transduction in these brain regions. Images shown in Figure 5 and the associated supplemental figure are maximum intensity projections.

### *Ex vivo* dendrite length, dendritic complexity, spine density and spine morphology analyses

Fixed tissue coronal slices of 50 um thickness were obtained from the Ptn x Thy1-YFP and Ts65Dn x Thy1-YFP lines. These preparations were used for the characterizations of total dendrite length, dendritic complexity, spine density and spine morphology of Layer 5 pyramidal neurons described in Figure 3/Figure S3, as well as the rescue experiments described in Figure 6/Figure S6. TFM was removed with a PBS wash and coverslips were mounted using Fluoromount G. Triangular pieces of parafilm were cut and placed at the top right and bottom left corners of the slides to prevent the coverslip from crushing the thick tissue and obscuring spine morphology. All subsequent analyses were performed blinded to genotype and treatment.

#### Imaging and analysis of total dendrite length and complexity

Imaging was performed on an LSM 700 confocal microscope. Six independent experiments were performed for Ptn WT vs. KO, Ts65Dn WT vs. Mut and the 4-group rescue experiments (WT smFP, WT Ptn, Ts65Dn Mut smFP and Ts65Dn Mut Ptn). Laser settings and gain were maintained constant within each experiment. Tiled z-stacks of the visual cortex were obtained to include at least 5 labeled cells per image. Images were obtained using a 20x objective. The z-steps were obtained at 0.5 um increments with a total of 60 steps. Three slices/per mouse were imaged for each experiment. Zen Blue 3.3 was used to stitch the images and the stitched z-stack images were then imported into Imaris where the filament tracing tool was used to manually trace individual dendrites. Dendrites were traced if they were completely contained in the z-stack and did not appear to be cut through the z-dimension. Imaris outputs total dendrite length as the sum of all individual processes for a given filament (dendrite). Sholl analysis was performed using 50 um increments from the cell body. Imaris outputs the number of branches intersecting with the Sholl circles as a function of distance from the cell body. Quantifications for total dendrite length and Sholl analysis were performed on both the average of all dendrites analyzed/mouse as well as the combined dendrites for each dataset (all dendrites analyzed from 6 mice combined) with the same results. Images shown in Figure 3 are maximum intensity projections. Filaments shown in Figure 6 were exported as TIFF files representing the scene from the Imaris analysis file.

#### Imaging and analysis of spine density and morphology

Imaging of dendritic spines was performed on a Zeiss LSM 880 confocal microscope with Airyscan imaging modality. Six independent experiments were performed for Ptn WT vs. KO and Ts65Dn WT vs. Mut. Eight independent experiments were performed for the 4-group rescue experiments (WT smFP, WT Ptn, Ts65Dn Mut smFP and Ts65Dn Mut Ptn). Laser settings and gain were maintained constant within each experiment. Images were obtained using a 63x objective with 2.0 zoom. The z-steps were obtained at 0.17 um increments and the number of steps for each image was determined on an individual dendrite basis to encompass the entire dendrite. Dendrites were chosen for imaging based on the following criteria: 1.) the cell body resided in Layer 5 and the dendrite reached to the upper layers (Layers 1-3), 2.) the primary branch point on the dendrite resided in the upper layers, and 3.) the dendrite in the image could be resolved from surrounding dendrite branches when focusing through the z-stack. For each individual mouse, five separate dendrites were imaged that were sampled from at least two separate tissue sections. The images were Airyscan processed in Zen and exported as TIFF files. These images were imported into NeuronStudio for spine density and morphology analysis. Spine heads were labeled within NeuronStudio and the spine head width and neck length measurements were exported. These measurements were used to assign spine morphology based on the classifications described previously ^30,31^. Quantification of spine density (number of spines/um) was performed on the average of analyzed dendrites/mouse and the combined total dendrites analyzed/dataset yielding the same results, with the exception that spine density in P30 Ts65Dn Mut mice was significantly reduced when combining all dendrites but not at the average of the biological replicate level (Figure 4 and Figure S4). Spine morphology quantifications were performed on the total of all dendrites analyzed per mouse and presented as the percentage of total spines for each morphological category. Images presented in Figure 3 and Figure 6 are maximum intensity projections.

#### Immunohistochemical synaptic staining, imaging and analysis

Fixed tissue coronal slices of 16 um thickness were obtained from the Ptn, Ts65Dn, and Ts65Dn x Thy1-YFP lines. These preparations were used for the characterizations of synaptic puncta described in Figure 4/Figure S4, as well as the rescue experiments described in Figure 7 (YFP negative mice from the Ts65Dn x Thy1-YFP lines). TFM was removed with a PBS wash. Slices were blocked and permeabilized in blocking buffer (10% goat serum in PBS with 0.2% Triton) for one hour at room temperature. Primary antibodies were diluted in antibody dilution solution (5% goat serum in PBS with 0.2% Triton) and stained overnight in a humidified chamber at 4C. The antibodies used for pre-synaptic puncta were Vglut1 (1:2000, guinea pig, Millipore AB5905) and Vglut2 (1:2000, guinea pig, Millipore AB2251-I). Post-synaptic puncta were labeled with GluA2 (1:400, rabbit, AB1768-I). The next day the slides were washed in PBS with 0.2% Triton three times for 5, 10 and 15 minutes. Appropriate secondary antibodies were used in antibody dilution solution. Secondary antibodies were applied for two hours at room temperature followed by three washes of 5, 10 and 15 minutes in PBS with 0.2% Triton. DAPI was applied (1:1000) for five minutes followed by a PBS wash. Coverslips were mounted with Fluoromount G. For the characterization experiments (Figure 4 and Figure S4) GluA2 was labeled on the 488-channel (1:500, goat anti-rabbit 488) and Vglut1/2 were labeled on the 594-channel (1:500, goat anti-guinea pig 594, Thermo Fisher Scientific A11076). For the rescue experiments, Vglut1 was labeled on the 488-channel (1:500, goat anti-guinea pig 488, Thermo Fisher Scientific A11073) and GluA2 was labeled on the 647-channel (1:500, goat anti-rabbit 647, Thermo Fisher Scientific A21245). In these stainings, anti-HA antibody (1:500, rat) was added to confirm viral transduction within each mouse and HA was labeled on the 555-channel (1:500, goat anti-rat). Experiments in each dataset were repeated over 5-8 independent experiments. Images were obtained using a 63x objective with 1.2 zoom on an LSM 880 microscope. The laser settings and gain were maintained constant within each experiment. Z-stacks of 0.4 um increments were obtained separately in Layer 1 for Vglut2/GluA2 and Layer 2/3 for Vglut1/GluA2 with a total of 10 z-steps per image. Three slices/per mouse were imaged for each experiment. Quantification of colocalized puncta was performed as previously described using the Imaris spots tool and the colocalization algorithm ^10,31,42^. Spot sizes were set to 0.4 um for Vglut1 and GluA2 puncta and 0.5 um for Vglut2 puncta with a colocalization distance of 0.7 um. Variability between individual staining experiments necessitated the data for Ptn KO or Ts65Dn Mut mice to be normalized to the WT within each experiment. Images shown in Figure 4 and Figure 7 are single z planes.

### Behavioral analysis

Behavioral testing was performed in the Salk Institute’s In Vivo Scientific Services (Animal Resources Department). Behavioral tests were assayed in the following order with 1-2 days rest in between each test: open field, spontaneous alternation Y maze and fear conditioning. All tests were performed in the morning between the hours of ZT1 to ZT5.5. Brain tissue was collected immediately after fear conditioning. All behavioral tests were performed blinded to genotype.

#### Open Field

Mice were habituated to the testing room for thirty minutes prior to initiation of the test. The room was lit by indirect lighting of ∼30 lux. Activity Monitor software (Med Associates) was used to assess the locomotor activity over a ten-minute period. Mice were placed into the open field chamber (43.2 x 43.2 x 30.5 cm; Med Associates Inc. ENV-515S-A) facing the outer plexiglass wall. Motion data was collected with Activity Monitor v5.92 software (Med Associates Inc.) and analyzed. At the conclusion of the test, mice were returned to their home cage and placed back into the housing facility. Analysis was performed using Activity Monitor v6.02 (Med Associates Inc.). Data was analyzed as 5-minute blocks across the ten-minute test duration. The total distance traveled between the 4 groups tested (WT smFP, WT Ptn, Ts65Dn Mut smFP and Ts65Dn Mut Ptn) was assessed.

#### Spontaneous Alternation Y Maze

Mice were habituated to the testing room for thirty minutes prior to testing. The room was lit by indirect lighting of ∼30 lux. The mouse was placed at the distal end of the starting arm (Arm A) and allowed to explore freely in any of the three arms (Arm A, Arm B, Arm C) over a testing period of eight minutes. The dimensions of arms were 17.0 cm x 6.5 cm x 23.0 cm. After the completion of the testing period the mouse was returned to its home cage. ANY-maze v7.2 software (San Diego Instruments) was used for motion tracking and automated analysis. The percentage of correct alternations was quantified as well as the total number of arm entries during the test duration. A correct alternation is defined as a triad of three entries without a repeat entry into a previous arm in the triad.

#### Fear Conditioning

Mice were habituated to the testing room for one hour prior to testing. The room was lit by indirect lighting of ∼30 lux. The fear conditioning protocol used in this study is a two-day protocol. On the first day, mice were placed into the conditioning chamber (24.0 x 30.0 x 21.0 cm, Med Associates Inc. VFC-008) for a 3-minute habituation period before being exposed to a series of foot shocks that are paired to an acoustic tone. The tone (90 dB/5000 Hz) is presented prior to each foot shock. The tones were presented at 180, 300 and 420 seconds for 30-second durations. The foot shocks were delivered directly after termination of the tone at 0.5 mA for two seconds. The mice learn to associate the foot shock with the tone and typically exhibit a freezing behavior. The mice were kept outside of the testing room in an anteroom to avoid pre-exposure to the acoustic tone. On the second day, the mice were tested in two separate paradigms. The first paradigm was used to test the contextual memory of the chamber. The mice were placed into the same chamber configuration as on the first day during the training period. The test duration was three minutes. In this paradigm there was no foot shock or tone and the mice exhibited freezing behavior simply in response to the contextual memory of the chamber. After the contextual memory paradigm, the mice were tested in a novel context and with the auditory cue and distinct visual, tactile and olfactory cues from the previous chamber. This paradigm tests the memory association of the auditory cue with the foot shock. The test duration was three minutes. The tone was presented three times at 20, 70 and 120 seconds for a duration of 30 seconds. Video Freeze software v.2.07.0.106 (Med Associates) was used for data analysis. Freezing was defined as a minimum motion threshold of 18 over 15 frames (0.5 seconds). The freezing behavior was quantified as the percentage of total time freezing within the test periods for the same context and novel context paradigms. For the learned association paradigm on the first day, the freezing behavior was quantified as the percentage of time freezing within defined periods (pre-shock, after first shock to second shock, second shock to third shock, and remainder of time after third shock).

#### Quantification and statistical analysis

Statistical analysis and data visualization was performed in Graphpad Prism 10. All tests were two-tailed. Data was tested for normality using either Shapiro-Wilk or Kolmogorov-Smirnov tests. For normally distributed data, T-tests (or one-sample t-tests) were used to compare two groups, one-way ANOVAs were used to compare three groups and two-way ANOVAs were used to compare multiple groups and treatments. If data was not normally distributed, pairwise comparisons were performed using Mann Whitney Rank Sum test (or one-sample Wilcoxon Signed Rank sum) and multiple comparisons were performed using Kruskal-Wallis on ranks. All bar graphs and line graphs represent mean ± SEM. Figures were generated using Adobe Illustrator 2022 and Biorender.

#### Additional resources

The data that support the findings of this study are available from the lead contact upon reasonable request.

## Notes

### Competing Interest Statement

The authors have declared no competing interest.

### Summary of Updates

This version of the manuscript has been updated in abstract, introduction, figures, supplemental files, discussion.

## References

1. Parker, S.E., Mai, C.T., Canfield, M.A., Rickard, R., Wang, Y., Meyer, R.E., Anderson, P., Mason, C.A., Collins, J.S., Kirby, R.S., and Correa, A. (2010). Updated National Birth Prevalence estimates for selected birth defects in the United States, 2004-2006. Birth Defects Res A Clin Mol Teratol 88, 1008–1016. 10.1002/bdra.20735.

2. Dierssen, M. (2012). Down syndrome: the brain in trisomic mode. Nature Reviews Neuroscience 13, 844–858. 10.1038/nrn3314.

3. Becker, L.E., Armstrong, D.L., and Chan, F. (1986). Dendritic atrophy in children with Down’s syndrome. Ann Neurol 20, 520–526. 10.1002/ana.410200413.

4. Marin-Padilla, M. (1976). Pyramidal cell abnormalities in the motor cortex of a child with Down’s syndrome. A Golgi study. J Comp Neurol 167, 63–81. 10.1002/cne.901670105.

5. Suetsugu, M., and Mehraein, P. (1980). Spine distribution along the apical dendrites of the pyramidal neurons in Down’s syndrome. A quantitative Golgi study. Acta Neuropathol 50, 207–210. 10.1007/bf00688755.

6. Takashima, S., Becker, L.E., Armstrong, D.L., and Chan, F. (1981). Abnormal neuronal development in the visual cortex of the human fetus and infant with down’s syndrome. A quantitative and qualitative Golgi study. Brain Res 225, 1–21. 10.1016/0006-8993(81)90314-0.

7. Clarke, L.E., and Barres, B.A. (2013). Emerging roles of astrocytes in neural circuit development. Nature Reviews Neuroscience 14, 311–321. 10.1038/nrn3484.

8. Chung, W.S., Allen, N.J., and Eroglu, C. (2015). Astrocytes Control Synapse Formation, Function, and Elimination. Cold Spring Harb Perspect Biol 7, a020370. 10.1101/cshperspect.a020370.

9. Farhy-Tselnicker, I., and Allen, N.J. (2018). Astrocytes, neurons, synapses: a tripartite view on cortical circuit development. Neural Dev 13, 7. 10.1186/s13064-018-0104-y.

10. Blanco-Suarez, E., Liu, T.F., Kopelevich, A., and Allen, N.J. (2018). Astrocyte-Secreted Chordin-like 1 Drives Synapse Maturation and Limits Plasticity by Increasing Synaptic GluA2 AMPA Receptors. Neuron 100, 1116–1132.e1113. 10.1016/j.neuron.2018.09.043.

11. Baldwin, K.T., and Eroglu, C. (2017). Molecular mechanisms of astrocyte-induced synaptogenesis. Curr Opin Neurobiol 45, 113–120. 10.1016/j.conb.2017.05.006.

12. Garcia, O., Torres, M., Helguera, P., Coskun, P., and Busciglio, J. (2010). A role for thrombospondin-1 deficits in astrocyte-mediated spine and synaptic pathology in Down’s syndrome. PLoS One 5, e14200. 10.1371/journal.pone.0014200.

13. 13. Araujo, B.H.S., Kaid, C., De Souza, J.S., Gomes da Silva, S., Goulart, E., Caires, L.C.J., Musso, C.M., Torres, L.B., Ferrasa, A., Herai, R., et al. (2018). Down Syndrome iPSC-Derived Astrocytes Impair Neuronal Synaptogenesis and the mTOR Pathway In Vitro. Mol Neurobiol 55, 5962–5975. 10.1007/s12035-017-0818-6.

14. Christopherson, K.S., Ullian, E.M., Stokes, C.C., Mullowney, C.E., Hell, J.W., Agah, A., Lawler, J., Mosher, D.F., Bornstein, P., and Barres, B.A. (2005). Thrombospondins are astrocyte-secreted proteins that promote CNS synaptogenesis. Cell 120, 421–433.

15. Chen, C., Jiang, P., Xue, H., Peterson, S.E., Tran, H.T., McCann, A.E., Parast, M.M., Li, S., Pleasure, D.E., Laurent, L.C., et al. (2014). Role of astroglia in Down’s syndrome revealed by patient-derived human-induced pluripotent stem cells. Nat Commun 5, 4430. 10.1038/ncomms5430.

16. Caldwell, A.L.M., Sancho, L., Deng, J., Bosworth, A., Miglietta, A., Diedrich, J.K., Shokhirev, M.N., and Allen, N.J. (2022). Aberrant astrocyte protein secretion contributes to altered neuronal development in multiple models of neurodevelopmental disorders. Nat Neurosci 25, 1163–1178. 10.1038/s41593-022-01150-1.

17. Cahoy, J.D., Emery, B., Kaushal, A., Foo, L.C., Zamanian, J.L., Christopherson, K.S., Xing, Y., Lubischer, J.L., Krieg, P.A., Krupenko, S.A., Thompson, W.J., and Barres, B.A. (2008). A transcriptome database for astrocytes, neurons, and oligodendrocytes: a new resource for understanding brain development and function. J Neurosci 28, 264–278. 10.1523/jneurosci.4178-07.2008.

18. Paveliev, M., Fenrich, K.K., Kislin, M., Kuja-Panula, J., Kulesskiy, E., Varjosalo, M., Kajander, T., Mugantseva, E., Ahonen-Bishopp, A., Khiroug, L., et al. (2016). HB-GAM (pleiotrophin) reverses inhibition of neural regeneration by the CNS extracellular matrix. Sci Rep 6, 33916. 10.1038/srep33916.

19. Kulesskaya, N., Molotkov, D., Sliepen, S., Mugantseva, E., Garcia Horsman, A., Paveliev, M., and Rauvala, H. (2021). Heparin-Binding Growth-Associated Molecule (Pleiotrophin) Affects Sensory Signaling and Selected Motor Functions in Mouse Model of Anatomically Incomplete Cervical Spinal Cord Injury. Front Neurol 12, 738800. 10.3389/fneur.2021.738800.

20. Vicente-Rodríguez, M., Gramage, E., Herradón, G., and Pérez-García, C. (2013). Phosphoproteomic analysis of the striatum from pleiotrophin knockout and midkine knockout mice treated with cocaine reveals regulation of oxidative stress-related proteins potentially underlying cocaine-induced neurotoxicity and neurodegeneration. Toxicology 314, 166–173. 10.1016/j.tox.2013.09.014.

21. Tang, C., Wang, M., Wang, P., Wang, L., Wu, Q., and Guo, W. (2019). Neural Stem Cells Behave as a Functional Niche for the Maturation of Newborn Neurons through the Secretion of PTN. Neuron 101, 32–44.e36. 10.1016/j.neuron.2018.10.051.

22. 22. Richards, S.E.V., Moore, A.R., Nam, A.Y., Saxena, S., Paradis, S., and Van Hooser, S.D. (2020). Experience-Dependent Development of Dendritic Arbors in Mouse Visual Cortex. J Neurosci 40, 6536–6556. 10.1523/jneurosci.2910-19.2020.

23. Blue, M.E., and Parnavelas, J.G. (1983). The formation and maturation of synapses in the visual cortex of the rat. II. Quantitative analysis. J Neurocytol 12, 697–712. 10.1007/bf01181531.

24. Li, M., Cui, Z., Niu, Y., Liu, B., Fan, W., Yu, D., and Deng, J. (2010). Synaptogenesis in the developing mouse visual cortex. Brain Res Bull 81, 107–113. 10.1016/j.brainresbull.2009.08.028.

25. Brill, J., and Huguenard, J.R. (2008). Sequential changes in AMPA receptor targeting in the developing neocortical excitatory circuit. J Neurosci 28, 13918–13928. 10.1523/jneurosci.3229-08.2008.

26. Farhy-Tselnicker, I., Boisvert, M.M., Liu, H., Dowling, C., Erikson, G.A., Blanco-Suarez, E., Farhy, C., Shokhirev, M.N., Ecker, J.R., and Allen, N.J. (2021). Activity-dependent modulation of synapse-regulating genes in astrocytes. Elife 10. 10.7554/eLife.70514.

27. Amet, L.E., Lauri, S.E., Hienola, A., Croll, S.D., Lu, Y., Levorse, J.M., Prabhakaran, B., Taira, T., Rauvala, H., and Vogt, T.F. (2001). Enhanced hippocampal long-term potentiation in mice lacking heparin-binding growth-associated molecule. Mol Cell Neurosci 17, 1014–1024. 10.1006/mcne.2001.0998.

28. Brunjes, P.C., and Osterberg, S.K. (2015). Developmental Markers Expressed in Neocortical Layers Are Differentially Exhibited in Olfactory Cortex. PLoS One 10, e0138541. 10.1371/journal.pone.0138541.

29. Feng, G., Mellor, R.H., Bernstein, M., Keller-Peck, C., Nguyen, Q.T., Wallace, M., Nerbonne, J.M., Lichtman, J.W., and Sanes, J.R. (2000). Imaging neuronal subsets in transgenic mice expressing multiple spectral variants of GFP. Neuron 28, 41–51. 10.1016/s0896-6273(00)00084-2.

30. Risher, W.C., Ustunkaya, T., Singh Alvarado, J., and Eroglu, C. (2014). Rapid Golgi analysis method for efficient and unbiased classification of dendritic spines. PLoS One 9, e107591. 10.1371/journal.pone.0107591.

31. Bosworth, A.P., Contreras, M., Novak, S.W., Sancho, L., Salas, I.H., Manor, U., and Allen, N.J. (2023). Astrocyte glypican 5 regulates synapse maturation and stabilization. bioRxiv, 2023.2003.2002.529949. 10.1101/2023.03.02.529949.

32. Viswanathan, S., Williams, M.E., Bloss, E.B., Stasevich, T.J., Speer, C.M., Nern, A., Pfeiffer, B.D., Hooks, B.M., Li, W.P., English, B.P., et al. (2015). High-performance probes for light and electron microscopy. Nat Methods 12, 568–576. 10.1038/nmeth.3365.

33. Kleschevnikov, A.M., Belichenko, P.V., Faizi, M., Jacobs, L.F., Htun, K., Shamloo, M., and Mobley, W.C. (2012). Deficits in cognition and synaptic plasticity in a mouse model of Down syndrome ameliorated by GABAB receptor antagonists. J Neurosci 32, 9217–9227. 10.1523/jneurosci.1673-12.2012.

34. Bao, X., Mikami, T., Yamada, S., Faissner, A., Muramatsu, T., and Sugahara, K. (2005). Heparin-binding growth factor, pleiotrophin, mediates neuritogenic activity of embryonic pig brain-derived chondroitin sulfate/dermatan sulfate hybrid chains. J Biol Chem 280, 9180–9191. 10.1074/jbc.M413423200.

35. Zeiss, C.J. (2021). Comparative Milestones in Rodent and Human Postnatal Central Nervous System Development. Toxicol Pathol 49, 1368–1373. 10.1177/01926233211046933.

36. Chini, M., and Hanganu-Opatz, I.L. (2021). Prefrontal Cortex Development in Health and Disease: Lessons from Rodents and Humans. Trends Neurosci 44, 227–240. 10.1016/j.tins.2020.10.017.

37. Au, H.K.E., Isalan, M., and Mielcarek, M. (2021). Gene Therapy Advances: A Meta-Analysis of AAV Usage in Clinical Settings. Front Med (Lausanne) 8, 809118. 10.3389/fmed.2021.809118.

38. Nahmani, M., and Erisir, A. (2005). VGluT2 immunochemistry identifies thalamocortical terminals in layer 4 of adult and developing visual cortex. J Comp Neurol 484, 458–473. 10.1002/cne.20505.

39. Kadomatsu, K., and Muramatsu, T. (2004). Midkine and pleiotrophin in neural development and cancer. Cancer Lett 204, 127–143. 10.1016/s0304-3835(03)00450-6.

40. Kucukdereli, H., Allen, N.J., Lee, A.T., Feng, A., Ozlu, M.I., Conatser, L.M., Chakraborty, C., Workman, G., Weaver, M., Sage, E.H., Barres, B.A., and Eroglu, C. (2011). Control of excitatory CNS synaptogenesis by astrocyte-secreted proteins Hevin and SPARC. Proc Natl Acad Sci U S A 108, E440–449. 10.1073/pnas.1104977108.

41. Allen, N.J., Bennett, M.L., Foo, L.C., Wang, G.X., Chakraborty, C., Smith, S.J., and Barres, B.A. (2012). Astrocyte glypicans 4 and 6 promote formation of excitatory synapses via GluA1 AMPA receptors. Nature 486, 410–414. 10.1038/nature11059.

42. 42. Farhy-Tselnicker, I., van Casteren, A.C.M., Lee, A., Chang, V.T., Aricescu, A.R., and Allen, N.J. (2017). Astrocyte-Secreted Glypican 4 Regulates Release of Neuronal Pentraxin 1 from Axons to Induce Functional Synapse Formation. Neuron 96, 428–445.e413. 10.1016/j.neuron.2017.09.053.

43. Takeda, A., Onodera, H., Sugimoto, A., Itoyama, Y., Kogure, K., Rauvala, H., and Shibahara, S. (1995). Induction of heparin-binding growth-associated molecule expression in reactive astrocytes following hippocampal neuronal injury. Neuroscience 68, 57–64. 10.1016/0306-4522(95)00110-5.

44. Davisson, M.T., Schmidt, C., Reeves, R.H., Irving, N.G., Akeson, E.C., Harris, B.S., and Bronson, R.T. (1993). Segmental trisomy as a mouse model for Down syndrome. Prog Clin Biol Res 384, 117–133.

45. Chan, K.Y., Jang, M.J., Yoo, B.B., Greenbaum, A., Ravi, N., Wu, W.L., Sánchez-Guardado, L., Lois, C., Mazmanian, S.K., Deverman, B.E., and Gradinaru, V. (2017). Engineered AAVs for efficient noninvasive gene delivery to the central and peripheral nervous systems. Nat Neurosci 20, 1172–1179. 10.1038/nn.4593.

