## Supplemental Figures with legends for "Dysregulation of astrocyte-secreted pleiotrophin contributes to neuronal structural and functional deficits in Down Syndrome"

Figure S1

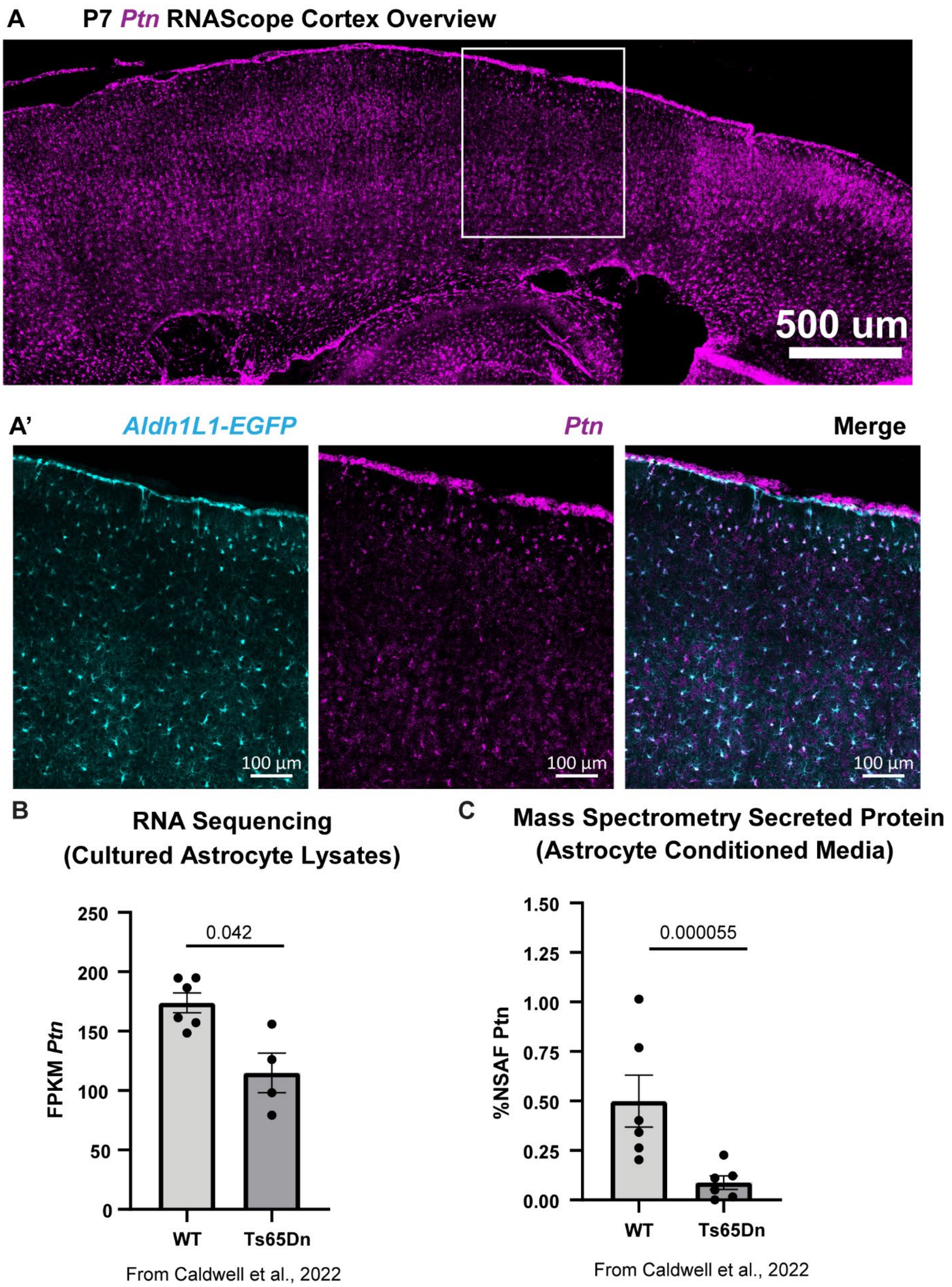


**Figure S1. *Ptn* mRNA is widespread throughout the cortex in neurotypical development and *Ptn* mRNA and secreted protein is reduced in DS astrocytes**

(A) Overview image of cortical *Ptn* (magenta) expression at P7. Scale bar = 500 um. (A') Zoomed image shows prominent *Ptn* in upper layer astrocytes (cyan). Scale bar = 100 um. (B) RNA-Sequencing of cultured primary cortical astrocytes shows reduced *Ptn* expression in Ts65Dn Mut astrocytes. Data from Caldwell et al., 2022. N = 4-6 cultures/group. (C) Mass spectrometry data of ACM shows reduced secreted Ptn protein from Ts65Dn Mut astrocytes. Data from Caldwell et al., 2022. N = 6 cultures/group. (B-C) Bar graphs represent mean ± SEM. Dots are each independent culture.

Figure S2

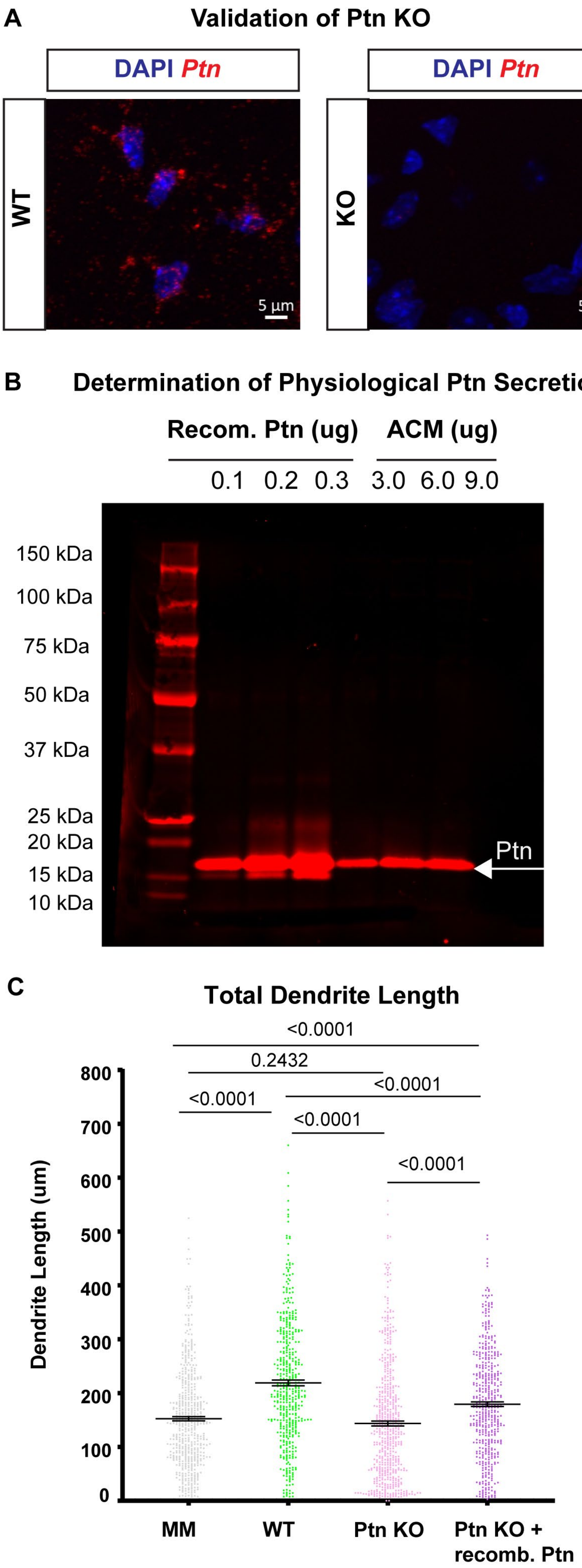
A
Validation of Ptn KO
DAPI Ptn
WT
KO
5 µm
5 µm
B
Determination of Physiological Ptn Secretion
Recom. Ptn (ug)
ACM (ug)
0.1 0.2 0.3 3.0 6.0 9.0
150 kDa
100 kDa
75 kDa
50 kDa
37 kDa
25 kDa
20 kDa
15 kDa
10 kDa
Ptn
C
Total Dendrite Length
<0.0001
0.2432
<0.0001
<0.0001
<0.0001
<0.0001
Dendrite Length (um)
800
700
600
500
400
300
200
100
0
MM
WT
Ptn KO
Ptn KO + recomb. Ptn

**Figure S2. Validation of Ptn KO, determination of physiological Ptn secretion and distribution data for *in vitro* dendrite outgrowth.**

(A) smFISH confirmed loss of *Ptn* mRNA in the visual cortex of Ptn KO mice at P30. Scale bar = 5 um. (B) Concentrated ACM from astrocyte cultures (3.0, 6.0 and 9.0 ug total) was run on a western blot and probed with an antibody against Ptn and the band intensity was compared to those of known recombinant Ptn protein input (0.1, 0.2 and 0.3 ug) to determine physiological levels of astrocyte Ptn secretion *in vitro*. Blot shown is representative of 4 independent experiments. (C) Distributions of dendrite length on the totality of cells analyzed from the 4 independent culture experiments described in Figure 2. Error bars represent mean ± SEM. Dots are individual cells. N = 460-566 cells/group. Statistics by Kruskal-Wallis test.

Figure S3

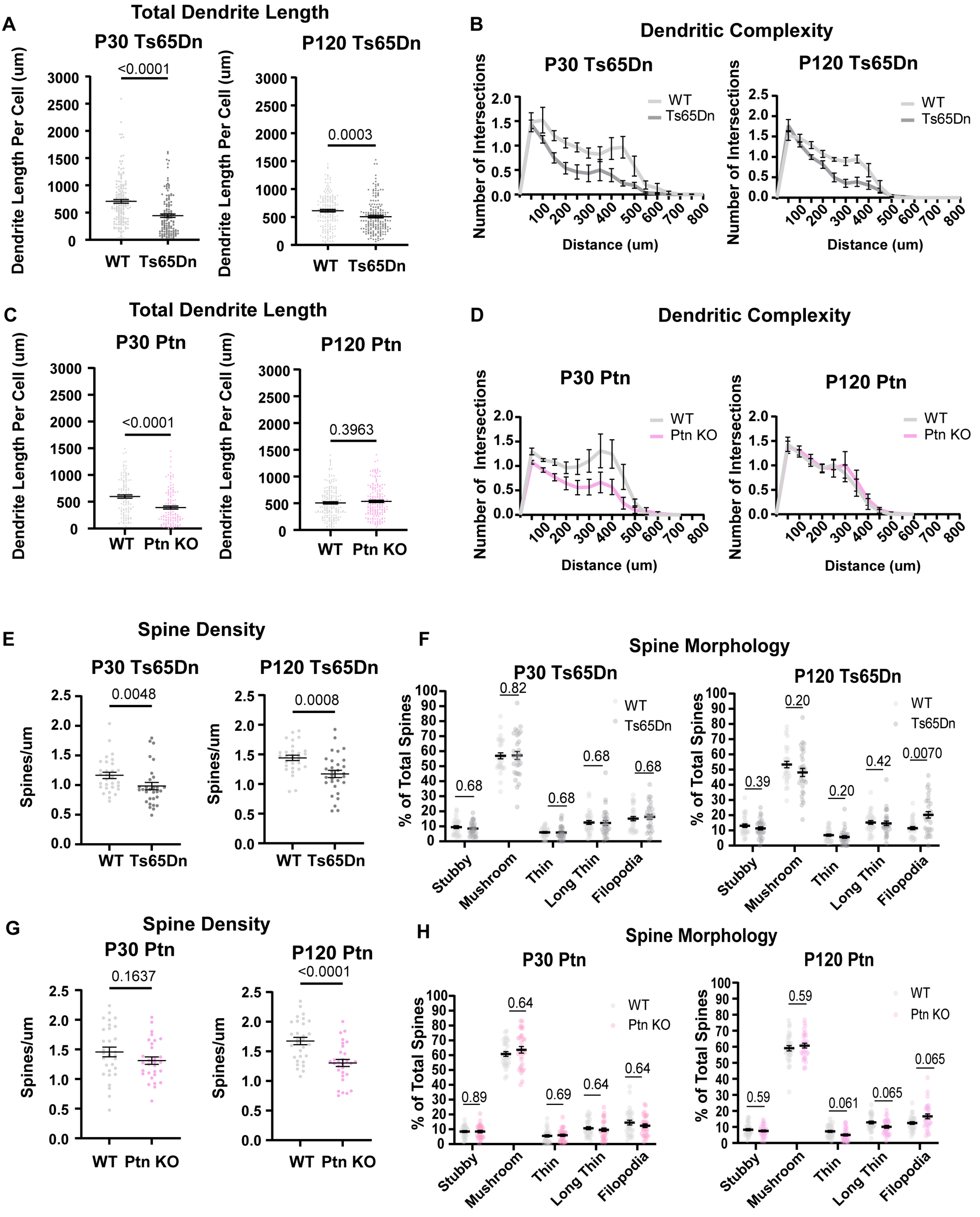
A
Total Dendrite Length
P30 Ts65Dn
P120 Ts65Dn
Dendrite Length Per Cell (um)
<0.0001
0.0003
WT
Ts65Dn
B
Dendritic Complexity
P30 Ts65Dn
P120 Ts65Dn
Number of Intersections
Distance (um)
WT
Ts65Dn
C
Total Dendrite Length
P30 Ptn
P120 Ptn
<0.0001
0.3963
WT
Ptn KO
D
Dendritic Complexity
P30 Ptn
P120 Ptn
Number of Intersections
Distance (um)
WT
Ptn KO
E
Spine Density
P30 Ts65Dn
P120 Ts65Dn
Spines/um
0.0048
0.0008
WT
Ts65Dn
F
Spine Morphology
P30 Ts65Dn
P120 Ts65Dn
% of Total Spines
0.68
0.82
0.68
0.68
0.68
0.39
0.20
0.20
0.42
0.0070
WT
Ts65Dn
Stubby
Mushroom
Thin
Long Thin
Filopodia
G
Spine Density
P30 Ptn
P120 Ptn
Spines/um
0.1637
<0.0001
WT
Ptn KO
H
Spine Morphology
P30 Ptn
P120 Ptn
% of Total Spines
0.89
0.64
0.69
0.64
0.64
0.59
0.59
0.061
0.065
0.065
WT
Ptn KO
Stubby
Mushroom
Thin
Long Thin
Filopodia

**Figure S3. Distribution data in the Ts65Dn and Ptn KO models for dendrite length, Sholl analysis, spine density and spine morphology**

(A) Distributions of individual dendrite lengths at P30 and P120 in WT and Ts65Dn Mut mice. n = 134-178 dendrites/group from six individual mice. Statistics by Mann-Whitney test. (B) Sholl curves for dendritic complexity at P30 and P120 in the Ts65Dn line show a decreased number of branch points at distances >100 um from the cell body. (C) Distributions of dendrite length for individual dendrites at P30 and P120 in Ptn WT and KO mice. The KO mice have decreased dendrite length at P30 that recovers by P120. n = 123-180 dendrites/group from six individual mice. Statistics by Mann-Whitney. (D) Sholl curves for P30 and P120 dendrite branching data in Ptn WT and KO mice show decreased branching at distances >100 um from the cell body at P30 but no difference at P120 in KO mice. (E) Distributions of spine density on individual dendrites from WT and Ts65Dn Mut mice at P30 and P120 show decreased spine density in Mut mice at both ages. n = 30 dendrites/group from six individual mice. Statistics by Mann-Whitney test (P30) or unpaired t-test (P120). (F) Distributions of spine morphology on individual dendrites show that Ts65Dn Mut mice retain a higher percentage of filopodia spines at P120. Statistics by multiple Mann-Whitney t-tests with Benjamini, Krieger and Yekutieli correction. (G) Distributions of spine densities on individual dendrites for Ptn WT and KO mice show that KO mice have decreased spine density at P120. n = 30 dendrites/group from six individual mice. Statistics by unpaired t-test. (H) Distributions for spine categorizations on individual dendrites in the Ptn WT and KO mice show no differences were observed. Statistics by multiple Mann-Whitney t-tests with Benjamini, Krieger and Yekutieli correction. Error bars represent mean ± SEM. Dots are individual dendrites.

Figure S4

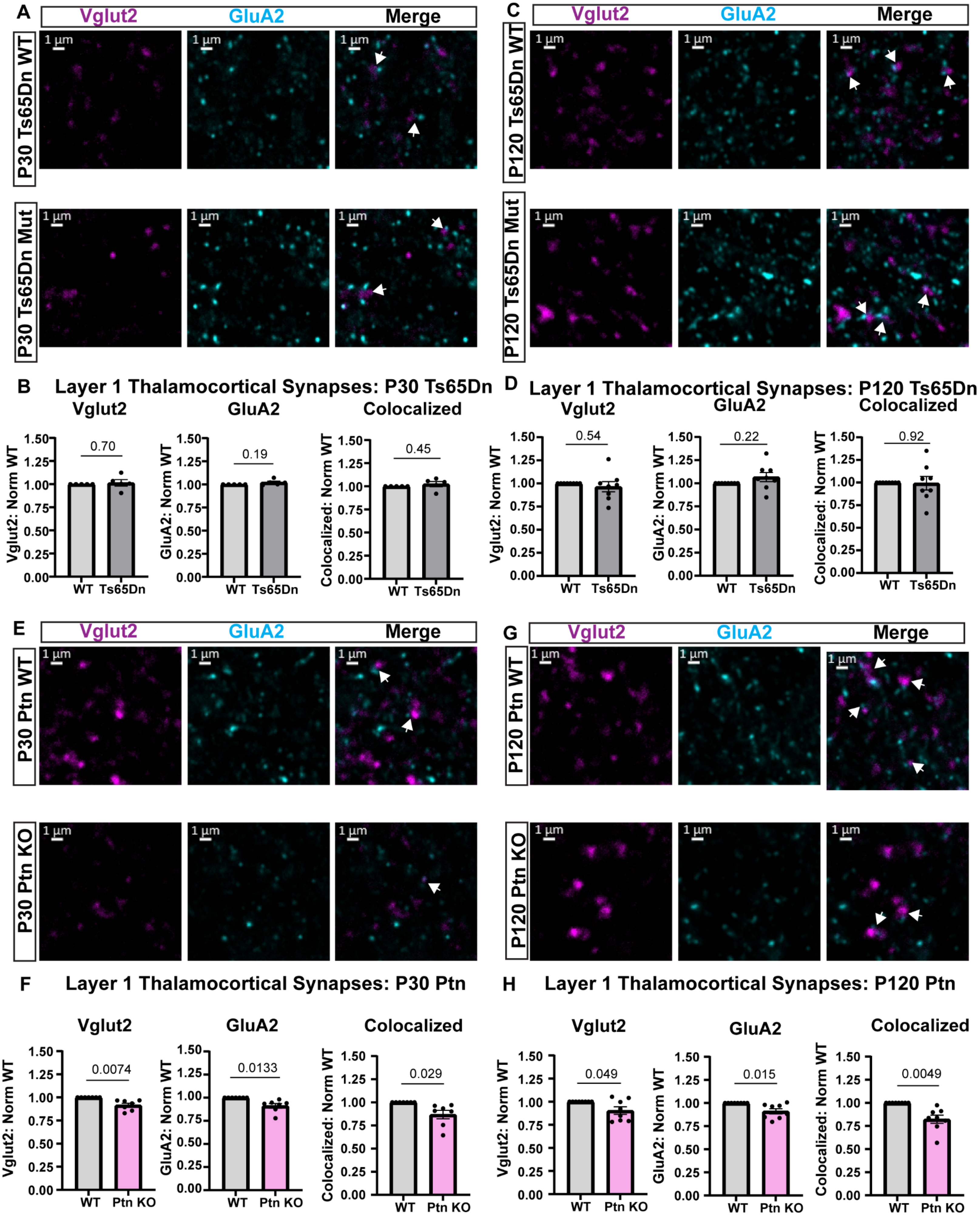
A
Vglut2
GluA2
Merge
P30 Ts65Dn WT
P30 Ts65Dn Mut
1 µm
C
P120 Ts65Dn WT
P120 Ts65Dn Mut
B
Layer 1 Thalamocortical Synapses: P30 Ts65Dn
Vglut2: Norm WT
GluA2: Norm WT
Colocalized: Norm WT
0.70
0.19
0.45
WT
Ts65Dn
D
Layer 1 Thalamocortical Synapses: P120 Ts65Dn
0.54
0.22
0.92
E
P30 Ptn WT
P30 Ptn KO
G
P120 Ptn WT
P120 Ptn KO
F
Layer 1 Thalamocortical Synapses: P30 Ptn
0.0074
0.0133
0.029
Ptn KO
H
Layer 1 Thalamocortical Synapses: P120 Ptn
0.049
0.015
0.0049
Colocalized

**Figure S4. Ptn KO mice have thalamocortical synapse phenotypes**

(A) Example images of thalamocortical pre-synaptic Vglut2 puncta (magenta), post-synaptic GluA2 puncta (cyan) and their colocalization in Ts65Dn WT and Mut mice at P30 in Layer 1. (B) Ts65Dn Mut mice do not show changes in thalamocortical synapses at P30. (C) Example images as in (A) at P120. (D) Ts65Dn Mut do not have thalamocortical synapse phenotypes at P120. (E) Representative images of Vglut2 (magenta), GluA2 (cyan) and their colocalization in Ptn WT and KO mice at P30. (F) Ptn KO mice have decreased levels of Vglut2, GluA2 and their colocalization compared to WT at P30. (G) Example images as in (E) at P120 in the Ptn line. (H) Decreased levels of Vglut2, GluA2 and their colocalization remains in Ptn KO mice at P120. Scale bars for all images = 1 um. N = 6-8 mice/group. Error bars show mean ± SEM. Dots are individual mice. Statistics by one-sample t-test.

Figure S5

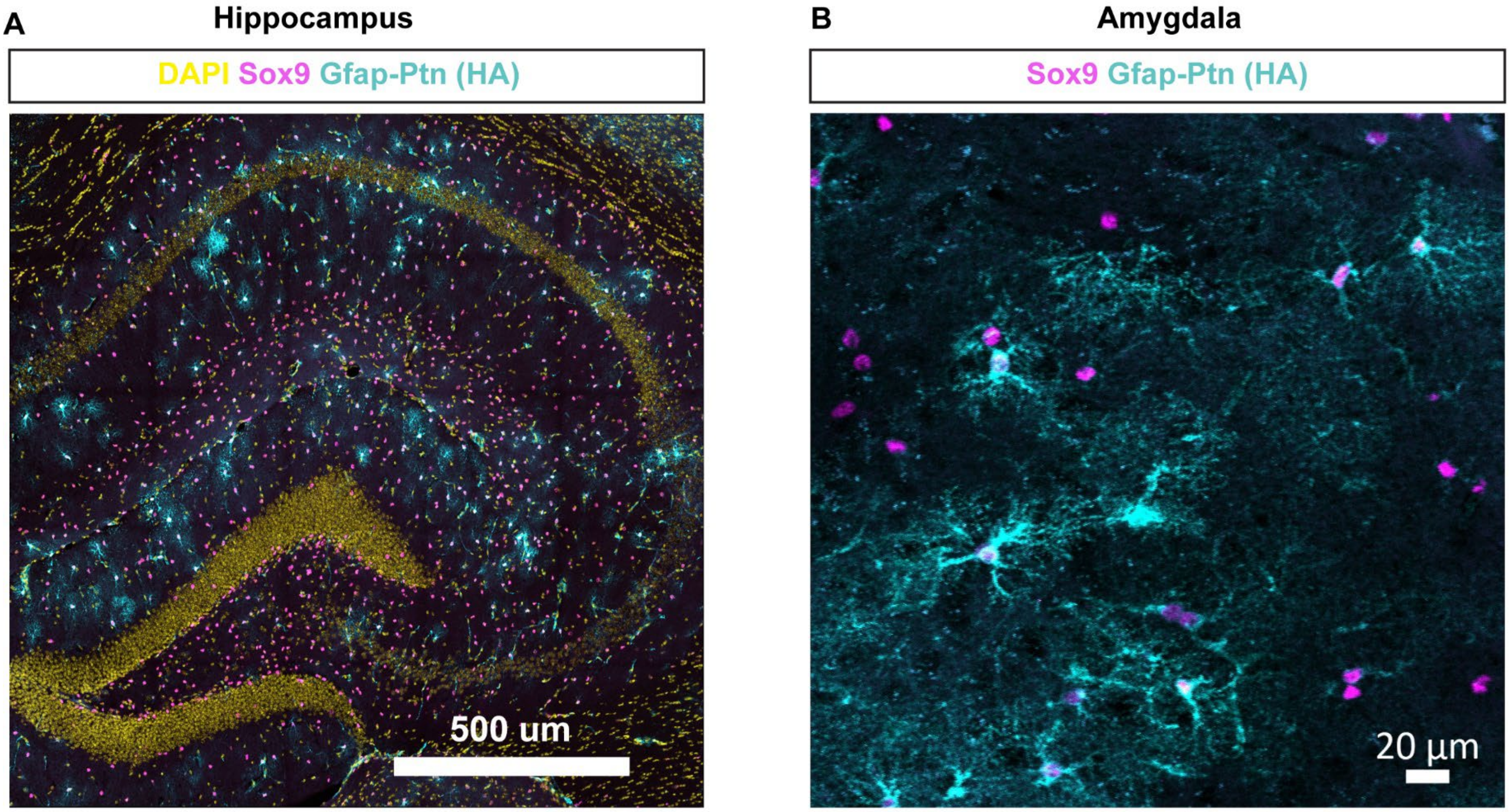


**Figure S5. Systemic viral injection strategy targets multiple brain regions.**

(A) Example image showing expression of Gfap-Ptn (cyan) in Sox9+ astrocytes (magenta) in hippocampus. The neuronal layers marked by DAPI (yellow) do not co-label with anti-HA in Gfap-Ptn-transduced cells. Scale bar = 500 um. (B) Example images of Gfap-Ptn virus (cyan) expression in Sox9+ (magenta) astrocytes in the lateral amygdala. Scale bar = 20 um.

Figure S6

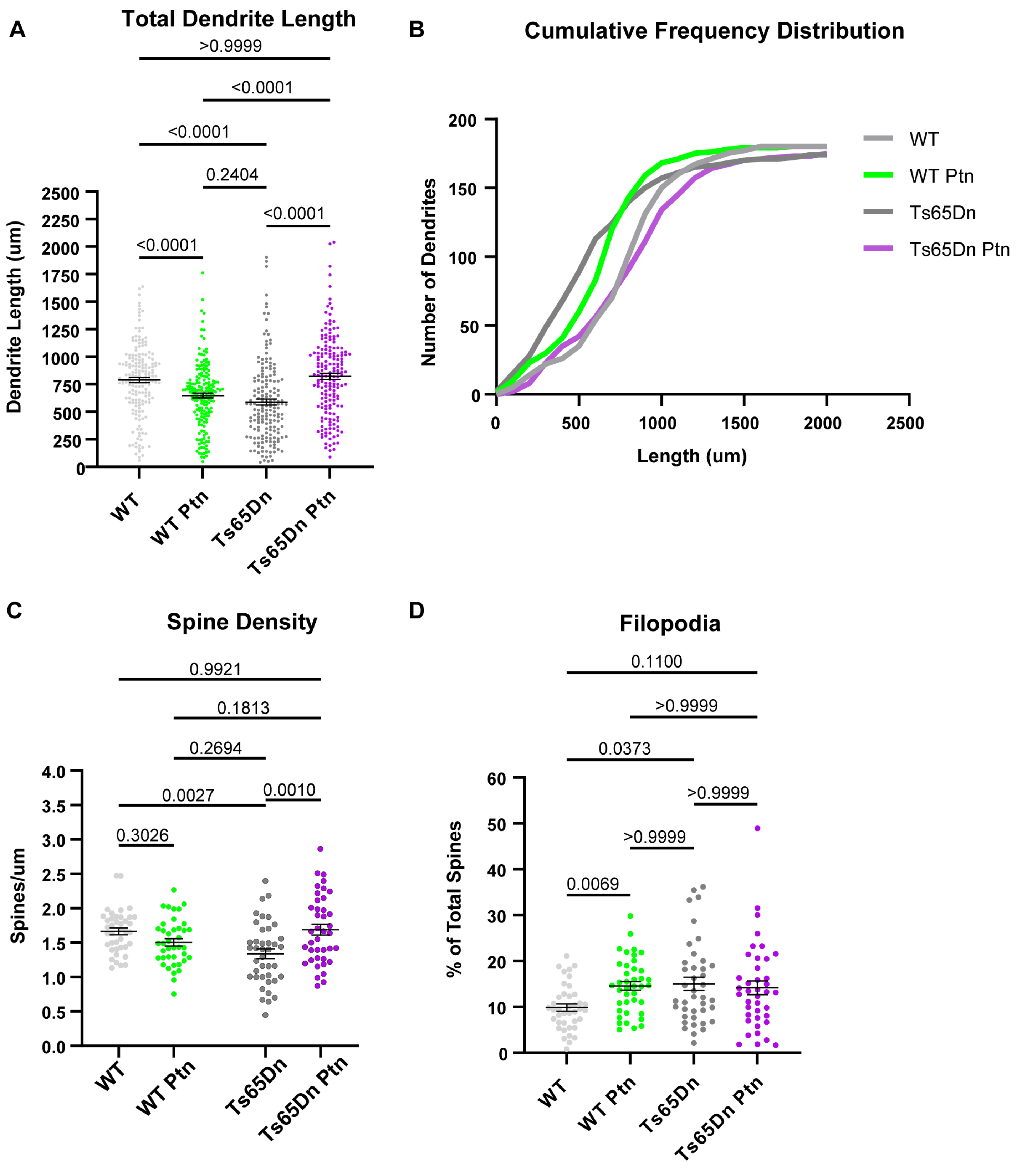


**Figure S6. Distribution data for total dendrite length and spine density**

(A) Distributions of total dendrite length for individual dendrites from 6 biological replicate (mice) groups. n = 174-180 dendrites/group. Statistics by Kruskal-Wallis with Dunn's test. (B) Cumulative frequency distribution curves for total dendrite length of individual dendrites in each group shows a shift towards decreased dendrite length in the Ts65Dn and WT Ptn groups. (C) Distributions of spine density for individual dendrites. n = 40 dendrites/group from 8 biological replicate (mice) groups. Statistics by two-way ANOVA with Tukey's test. (D) Quantification of the percentage of filipodia spines shows that Ts65Dn and WT Ptn dendrites have an increased proportion of filopodia spines compared to WT smFP. Statistics by Kruskal-Wallis with Dunn's test. Error bars represent mean ± SEM. Dots are individual dendrites.

Figure S7

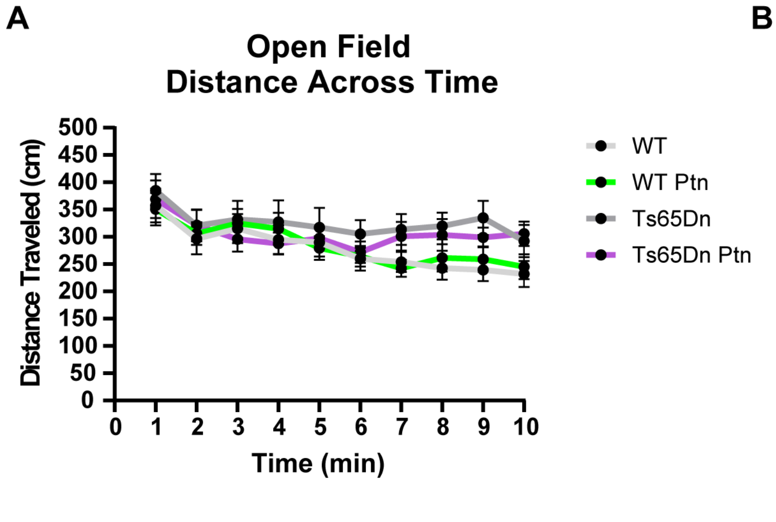


**Figure S7. Open field assessment of hyperactivity**

(A) The distance traveled in the open field across a 10-minute duration in 1-minute blocks is shown for each group. (B) Quantification of total distance traveled during the 1st 5-minute block shows no genotype difference. Statistics by Kruskal-Wallis with Dunn's. (C) Quantification of total distance traveled during the 2nd 5-minute block shows a main genotype effect of hyperactivity in Ts65Dn mice. Statistics by two-way ANOVA with Dunnett's. N = 18-24 mice/group. Error bars represent mean of each group ± SEM. Dots in B-C are individual mice.
